# ConnecTF: A platform to build gene networks by integrating transcription factor-target gene interactions

**DOI:** 10.1101/2020.07.07.191627

**Authors:** M.D. Brooks, C.-L. Juang, M.S. Katari, J.M. Alvarez, A.V. Pasquino, H.-J. Shih, J. Huang, C. Shanks, J. Cirrone, G.M. Coruzzi

**Affiliations:** Center for Genomics and Systems Biology, Department of Biology, New York University, NY, USA; Centro de Genómica y Bioinformática, Facultad de Ciencias, Universidad Mayor, Santiago, Chile; Millennium Institute for Integrative Biology (iBio), Santiago, Chile; Courant Institute for Mathematical Sciences, Department of Computer Science, New York University, NY, USA

## Abstract

Deciphering gene regulatory networks (GRNs) is both a promise and challenge of systems biology. The promise is identifying key transcription factors (TFs) that enable an organism to react to changes in its environment. The challenge is constructing GRNs that involve hundreds of TFs and hundreds of thousands of interactions with their genome-wide target genes validated by high-throughput sequencing. To address this challenge, we developed ConnecTF, a species-independent web-based platform for constructing validated GRNs and to refine inferred GRNs via combined analysis of genome-wide studies of TF-target gene binding, TF-target regulation and other TF-centric omic data. We demonstrate the functionality of ConnecTF in three case studies, showing how integration within and across TF-target datasets uncovers biological insights. Case study 1 uses integration of TF-target gene regulation and binding datasets to uncover mode-of-action and identify potential TF partners for 14 TFs in abscisic acid signaling. Case study 2 demonstrates how genome-wide TF-target data and automated functions in ConnecTF are used to conduct precision/recall analysis and pruning of an inferred GRN for nitrogen signaling. In case study 3, we use ConnecTF to chart a network path from NLP7, a master TF in nitrogen signaling, to direct secondary TF_2_s, to its indirect targets, in an approach called Network Walking. The public version of ConnecTF (https://ConnecTF.org) contains 3,738,278 TF-target interactions for 423 TFs in Arabidopsis, and 839,210 TF-target interactions for 139 TFs in maize. The database and tools in ConnecTF should advance the exploration of GRNs in plant systems biology applications for models and crops.

## Introduction

Deciphering gene regulatory networks (GRN) is an important task, as it can reveal regulatory loci, transcription factors (TFs), crucial for development, stress responses, or disease, with potential applications in agriculture and medicine (Petricka et al., 2012; Chatterjee and Ahituv, 2017; Gupta and Singh, 2019). However, integrating experimentally validated connections between TFs and their genome-wide target genes in such GRNs remains a challenge.

With the advent of next-generation sequencing, there are a growing number of methods to validate TF-target gene connections within GRNs, each with its own set of benefits and drawbacks. Methods that provide evidence for where a TF is likely to bind to the genome include; chromatin immunoprecipitation (ChIP-seq), DNA affinity purification sequencing (DAP-seq) (O’Malley et al., 2016), and cis-motif enrichment. To determine when TF-binding leads to target gene regulation requires the integration of TF-binding data with TF-regulation datasets. However, large-scale datasets that validate TF-target gene regulation data are sparse relative to TF-target gene binding data. This is largely due to the low throughput nature of TF-perturbation approaches in planta (e.g. overexpression or mutants). Thus, there is a need for higher throughput methods to rapidly identify direct regulated TF-targets in plants. One such method is the Transient Assay Reporting Genome-wide Effects of Transcription factors (TARGET) which uses temporal controlled TF nuclear entry to identify direct regulated TF-targets in isolated plant cells (Bargmann et al., 2013; Brooks et al., 2019).

Such large-scale datasets for TF-target binding or regulation can be used to verify predictions of TF-target gene connections in GRNs (Marbach et al., 2012; Banf and Rhee, 2017; Mochida et al., 2018; Kulkarni and Vandepoele, 2019). Validated TF-target interactions can also be used as priors (e.g. “ground truths”) to train machine learning in network inference methods (Greenfield et al., 2013; Petralia et al., 2015; Cirrone et al., 2020), and/or as a gold standard with which to benchmark/refine the accuracy of predicted TF-target interactions in learned GRNs (Marbach et al., 2012; Varala et al., 2018; Brooks et al., 2019). We have previously shown how the integration of TF-target binding with TF-target regulation datasets can reveal distinct modes-of-action of a TF on induced vs. repressed gene targets (Brooks et al., 2019).

Platforms that facilitate access to and integration of such large-scale datasets that validate TF-target gene interactions are crucial to accelerate studies of validated and inferred GRNs. To this end, there are efforts to aggregate TF-target datasets, largely TF-binding and cis-motif elements, for many species, including human (Han et al., 2018), yeast (Monteiro et al., 2019), *E. coli* (Santos-Zavaleta et al., 2019), and Arabidopsis (Yilmaz et al., 2010; Kulkarni et al., 2018; Tian et al., 2019). There are also web portals that provide access to specific experimental datasets that support TF-target binding, for example the Plant Cistrome database for large scale assays of in vitro TF-target binding (DAP-seq) (O’Malley et al., 2016). Primarily, these platforms allow users to query a TF and obtain a list of TF-bound target genes or vice versa.

Despite these advances, few, if any, platforms enable a combined analysis of TF-bound genes, TF-regulated genes, and co-expression data, or the ability to combine such datasets to refine/validate predicted GRNs. An important feature missing from most available web tools is the ability to integrate genome-wide targets of a single TF validated by different experimental approaches (e.g. ChIP-seq, DAP-Seq and RNA-seq), captured under the same or different experimental conditions. A second feature that is currently lacking is the ability to compare the validated targets of multiple TFs and determine their hierarchy in a GRN, as they relate to a set of user-defined genes such as a pathway of interest. Finally, tools are also needed to facilitate the refinement/pruning of predicted GRNs by using the validated TF-target interactions from genomic studies to perform precision/recall analysis.

To meet the need in the plant systems biology community to build, validate and refine GRNs, we developed ConnecTF, a platform which offers a query interface to access a TF-centric database consisting of large-scale validated TF-target gene interactions based on TF-target binding (e.g. ChIP/DAP-Seq) and other gene-to-gene directed (e.g. TF-target regulation,) or undirected (e.g. TF-TF protein-protein interaction) relationships. We are hosting a publicly available instance of ConnecTF (https://ConnecTF.org) which includes a database of large-scale validated TF-target interactions containing; TF-binding (in vivo and in vitro), TF-regulation (in planta and in plant cells), and cis-motif datasets for the model plant Arabidopsis and a crop, maize. The ConnecTF database currently contains 3,738,278 experimentally validated TF-target edges for 423 TFs in Arabidopsis (Table 1), and 839,210 experimentally validated TF-target edges for 139 TFs in maize (Supplemental Table 1). The database also includes the largest TF-target regulation dataset in plants, the direct regulated targets for 58 TFs in Arabidopsis (Varala et al., 2018; Brooks et al., 2019; Alvarez et al., 2020; this study)

**Table 1.** Overview of the validated Arabidopsis TF-target datasets in the ConnecTF database

| Interaction Type | Experiment Type | No. of TFs | # of edges | Reference |
| --- | --- | --- | --- | --- |
| TF-Binding | ChIP-seq | 26 | 257,400 | (Song et al., 2016)<br>(Birkenbihl et al., 2017) |
|  | DAP-seq | 382 | 3,335,595 | (O'Malley et al., 2016) |
| TF-Regulation | in planta perturbation | 3 | 7,894 | (Marchive et al., 2013)<br>(Varala et al., 2018) |
|  | TARGET (plant cells) | 58 | 137,389 | (Brooks et al., 2019)<br>(Alvarez et al., 2020)<br>(Brooks et al., 2020 this study) |
| TF-TF protein-protein interactions | HaloTag-NAPPA CrY2H | 1,221 | 6,555 | (Yazaki et al., 2016)<br>(Trigg et al., 2017) |
| Cis-binding motifs | TF cis-binding motifs | 1310 cis-motifs for 730 TFs collected from Cis-BP |  | (Weirauch et al., 2014) |
|  | Cis-motif clusters | 80 clusters from 1,282 individual cis-binding motifs |  | (Brooks et al., 2019) |

We demonstrate in three case studies how the features of ConnecTF and its ability to integrate a large and diverse variety of validated TF-target gene datasets can provide biological insights into GRNs. In the first case study, we demonstrate how the integration of validated TF-binding and TF-regulation datasets enabled us to discover how TFs and their TF-TF partner interactions influence the regulation of genes in the abscisic acid (ABA) pathway. In the second case study, we demonstrate how ConnecTF can be used to facilitate precision/recall analysis of inferred nitrogen regulatory networks using gold standard validated TF-target interactions stored in the ConnecTF database. In the third case study, we demonstrate how the query system of ConnecTF can be used to integrate validated TF-target datasets from multiple TFs into a unified network path. Specifically, using the query functions in ConnecTF, we were able to chart a network path from the direct targets of - NIN-LIKE PROTEIN 7 (NLP7), a key TF in the nitrogen response (Marchive et al., 2013; Alvarez et al., 2020), to its indirect targets in planta, using an adaptation of a Network Walking approach (Brooks et al., 2019). Overall, the database and analysis/integration tools of ConnecTF can be used to advance the validation of GRNs involved in any pathway using systems biology approaches in models or crops.

## Results

### ConnecTF: A query interface and database to integrate TF-target gene interactions of different data types

The ConnecTF platform and database enables researchers to access, analyze and integrate large-scale experimentally determined datasets on TF-target gene interactions including TF-binding, TF-regulation, TF-TF protein interactions, and cis-motifs (Table 1 and Supplemental Table 1). An important feature of ConnecTF is that it not only provides researchers access to the large-scale validated TF-target datasets housed in the database, but also offers a user-friendly interface to perform analyses to combine these various datasets for one or many TFs. This includes the ability for users to provide their own target gene lists or predicted networks and identify the TFs that regulate their pathway/network of interest. Users can also provide their own inferred networks and use the validated TF-target data in the ConnecTF database as a gold standard to perform precision/recall analysis using automated functions in ConnecTF. These applications are described in the three case studies below.

The backend structure and tools available in ConnecTF are species-independent and built using common software (Supplemental Figure 1). The source code and detailed instructions on how to setup a personal version of ConnecTF are available on GitHub (https://github.com/coruzzilab/connectf_server). This will enable others to setup their own instance of ConnecTF for public or private sharing of TF-centric genomic data. We are hosting public versions of ConnecTF with large-scale TF-target validation datasets from Arabidopsis (https://ConnecTF.org/) or maize (https://Maize.ConnecTF.org/). The current version of the Arabidopsis ConnecTF database primarily houses TF-binding or TF-regulation datasets that have been performed at scale (Table 1), enabling direct comparisons of TF-target interactions. This database includes; 388 Arabidopsis TFs for which TF-target binding was identified in vitro by DAP-seq (O’Malley et al., 2016), 21 TF-target binding datasets identified in planta by ChIP-seq (Song et al., 2016), and 58 TFs for which direct regulated TF-target genes were identified in isolated plant cells (Varala et al., 2018; Brooks et al., 2019; Alvarez et al., 2020), including 14 TFs from this study (Supplemental Table 3). For maize, the ConnecTF datasets include the recently reported ChIP-seq data for 103 TFs performed in isolated maize cells (Tu et al., 2020), TF perturbation and ChIP binding datasets collected from the literature (Bolduc et al., 2012; Morohashi et al., 2012; Eveland et al., 2014; Li et al., 2015), as well as in vitro TF-target binding identified by DAP-seq for 32 maize TFs (Ricci et al., 2019). In addition, for both Arabidopsis and maize, we have included in the database ATAC-seq (Lu et al., 2019) and DNA Hypersensitivity (DHS) (Sullivan et al., 2014) datasets, which enables users to filter TF-target interactions (e.g. TF-target gene binding) for those occurring in open chromatin regions of the different tissues from those studies.

A key feature of ConnecTF is its logic-based query system. A query in ConnecTF is built by constructing a series of constraints to restrict the set of TFs, the set of target genes, the type of interaction (e.g. TF-target edge type), or other attributes associated with the data. The result of the query is the network (or subnetwork) of interactions for the selected set of TFs and their targets. This query system allows users to select a single TF or multiple TFs of interest, filter the TF-targets based on different criteria (e.g. regulation by a signal of interest, e.g. ABA), and integrate validated TF-target data across multiple TFs. This includes the ability to search for targets of all TFs in the database, or a selected subset of TFs of interest. The query system also allows users to perform analyses based on the experimental type of validated TF-target interaction (e.g. TF-binding) or any other criteria in the metadata (e.g. TF-target assays performed in leaf vs. root). Queries can be built using the graphical *Query Builder* interface or by typing queries into the search text box. This makes the query system easy to use both for researchers new to the ConnecTF site, and for those who wish to build complex queries to parse multiple types of experimentally verified TF-target datasets for the TFs available in the database.

ConnecTF includes several analysis and visualization tools for data integration (Figure 1), whose utility we demonstrate in three case studies. Once a query has been submitted and is processed, the *Summary* tab is loaded and gives an overview of the total number of validated TF-target genes for each experiment that was queried, grouped by individual TFs. The validated TF-target interactions are then made available in the *Table* tab, which provides an interactive table that can be downloaded for offline use in either Excel or CSV formats. The five remaining tabs in ConnecTF, allow users to analyze the queried data in various ways (Figure 1): 1) *Network* tab – provides access to TF-target network as JSON or SIF files or visualized using Cytoscape,js (Franz et al., 2015) (Figure 1A), 2) *Target List Enrichment* tab – displays the overlap between user-submitted gene list(s) and validated TF-targets bound and/or regulated by the queried TF(s) and calculates statistical enrichment 3) *Motif Enrichment* tab – performs statistical tests for cis-motif enrichment of TF-binding sites in the validated targets of queried TFs (Figure 1E), 4) *Gene Set Enrichment* tab – calculates the significance of overlap between the validated targets of each TF analysis, when compared pairwise (Figure 1C), and 5) *Sungear* tab –compares the overlaps between TF-targets from multiple gene lists, comparable to a Venn diagram, but better suited to analyzing more than three lists (Figure 1D) (Poultney et al., 2006). The *Network* tab also enables users to upload a predicted network and use validated TF-target datasets housed in the ConnecTF database to perform an automated precision/recall analysis. This function generates an area under precision recall (AUPR) curve with an interactive sliding-window feature that can be used to select a precision cutoff with which to prune/refine the predicted network (Figure 1B) (Marchive et al., 2013; Banf and Rhee, 2017). The three case studies below provide use examples for ConnecTF by combining each of these features.

**Figure 1.**
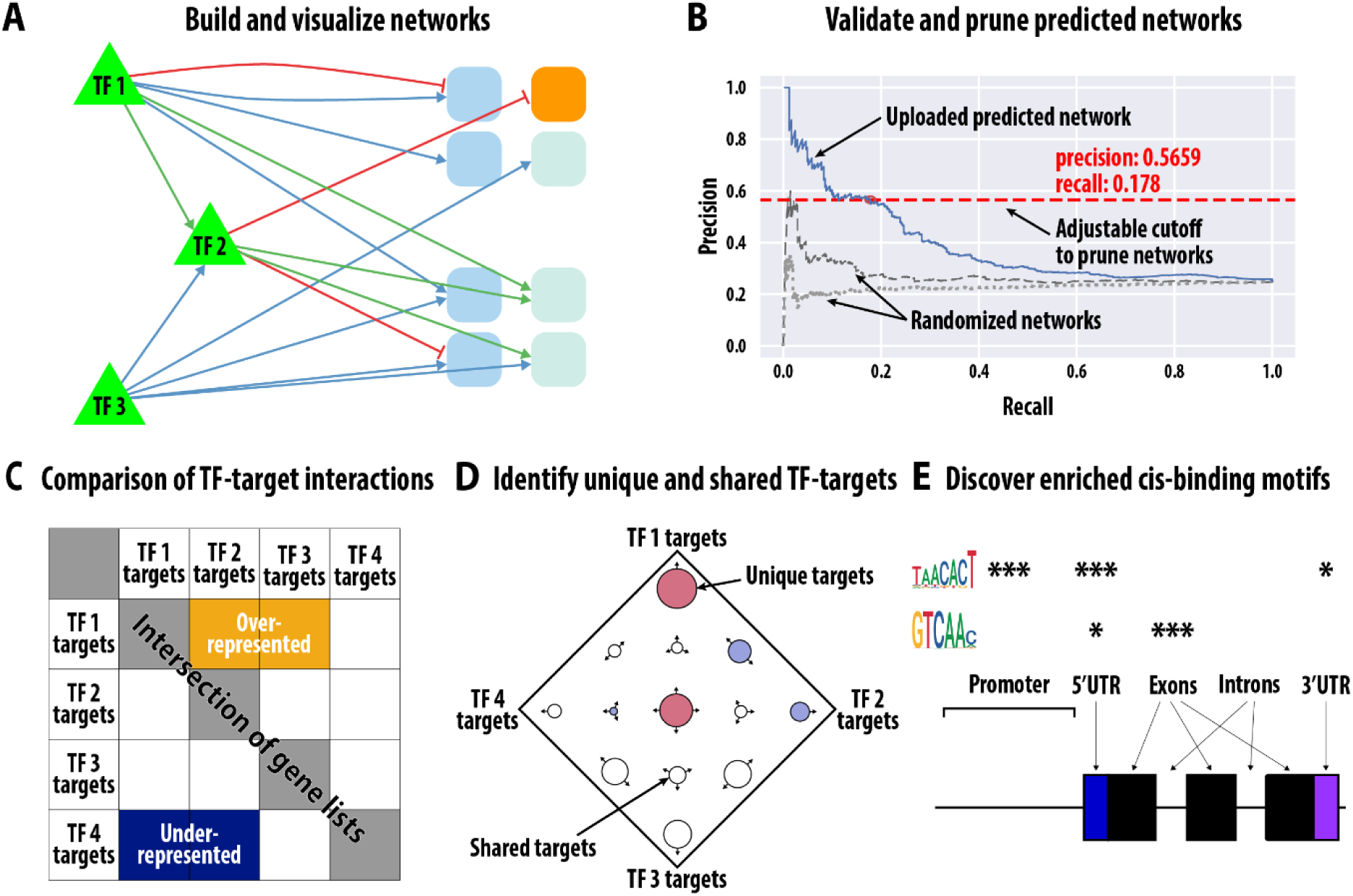
Analysis and visualization tools in ConnecTF for the integration of data supporting TF-target gene interactions to build/validate gene regulatory networks. ConnecTF contains TF-target interactions for 707 experiments from Arabidopsis and 158 experiments in maize for a total of 4.58 million TF-target interactions for 590 TFs (Table 1 and Supplemental Table 1). The distinct types of validated TF-target data within each species can be filtered and integrated using analysis/visualization tools within ConnecTF to; A) build and visualize validated gene regulatory networks, B) use validated TF-target data to perform precision/recall analysis and prune predicted networks (user uploaded or predefined in database), C) compare whether the TF-targets in common between two experiments/TFs are over-represented or under-represented, D) determine how TF-targets are distributed between TF experiments, and E) identify enriched cis-binding motifs in validated TF targets.

### Getting Started: Basic queries in ConnecTF

The most basic query in ConnecTF is to enter a TF name/symbol or Gene ID, which will return all of the experiments that validate TF-target gene interactions for that TF in the database. To demonstrate, we submitted a query for NLP7 (AT4G24020), a master regulator in the nitrogen signaling pathway, and the results returned from the ConnecTF database include seven experiments for NLP7: four ChIP-seq experiments performed in isolated root cells (Alvarez et al., 2020), one in vitro TF-target binding experiment using DAP-seq (O’Malley et al., 2016), one TF overexpression experiment that identifies direct regulated targets of NLP7 in isolated root cells (Alvarez et al., 2020), and one experiment identifying NLP7-regulated targets based on analysis of an *nlp7* mutant in planta (Marchive et al., 2013). These results can be viewed in the *Table* tab on the ConnecTF site or downloaded as an Excel file (Supplemental Table 2), and list the validated NLP7 target genes from any one of these experiments. This list includes descriptions of the validated NLP7 target genes (where available) and other details such as edge count (e.g. number of experiments where an interaction between the TF and this target are validated), *P*-value and log2 fold change, if available.

Determining the validated TF-target genes within a pathway or network of interest for one TF, or a set of TFs, is another common task that can be readily performed using ConnecTF. When a query is submitted in ConnecTF, the user can limit the target genes to one or more lists of genes using the *Target Gene List* box located below the *Query Builder*. We demonstrate this feature using the same NLP7 query as above, but in this example, from the *Target Gene List* box we select the predefined list of nitrogen response genes from shoot and root (Varala et al., 2018) named “Nitrogen_by_Time”. By selecting this list, the validated targets of NLP7 retrieved from the ConnecTF database are now restricted to the genes that are in one of these two pre-defined sets of genes (N-response in roots or shoots). In the results *Table* tab for this query, there are two additional columns that indicate each gene list (e.g. roots or shoots) to which the validated NLP7 targets belong (Supplemental Table 2). Uploading a *Target Gene List* also allows the user to determine the enrichment of gene targets of the TF in that pathway viewed in the *Target List Enrichment* tab.

### Case Study 1: Uncovering mechanisms of TF action and TF-TF interactions by integrating TF-target binding, TF-regulation and cis-element datasets

In this case study, we demonstrate how to use the query functions and data housed in ConnecTF to integrate TF-target gene regulation and TF-binding data to elucidate TF mode-of-action, including its potential TF partners. In our previous study of 33 TFs, we showed that a single TF can either induce *or* repress target genes (Brooks et al., 2019). Moreover, we showed examples where direct TF-target binding (e.g. via cis-motif enrichment and DAP-seq binding) was associated with TF-mediated target gene induction, while indirect binding via TF partner(s) (e.g. only captured by ChIP) could account for TF-mediated repression of a target gene (Brooks et al., 2019). However, we were unable to generalize this discovery, as only 3/33 TFs in that study had both vitro and in vivo TF-binding data. To expand and generalize our discoveries of these distinct TF modes-of-action, we used ConnecTF to integrate TF-regulation data (Supplemental Table 3), and TF-binding data (Song et al., 2016) for 14 TFs in the ABA signaling pathway. We did this by using functions in ConnecTF to integrate; i) the direct regulated TF targets of these 14 TFs identified in root cells (Supplemental Table 3) using the TARGET system (Bargmann et al., 2013; Brooks et al., 2019), ii) in planta TF-binding (e.g. ChIP-seq) (Song et al., 2016), iii) at least one cis-binding motif available on Cis-BP (Weirauch et al., 2014), and iv) validated in vitro TF-binding data obtained by DAP-seq (O’Malley et al., 2016) for 5/14 of the ABA responsive TFs.

#### Validated targets of 14 TFs are specifically enriched in ABA-responsive genes

First, we demonstrate how the validated TF-target gene datasets for these 14 ABA responsive TFs housed in the ConnecTF database can be integrated to understand how they regulate ABA signaling. To do this, we first used the *Target List Enrichment* tool in ConnecTF to determine for each of the 14 TFs whether the validated TF-regulated target genes identified by controlled TF-nuclear import in root cells using the TARGET assay (Bargmann et al., 2013; Brooks et al., 2019) were significantly enriched in a list of ABA responsive genes identified in Song et al. (Song et al., 2016). This integrated analysis showed that the direct regulated targets of these 14 TFs are each significantly enriched for ABA responsive genes (Fisher’s Exact test, *P*-value<0.05) (Figure 2, see Supplemental Data for query used to generate this figure). This analysis enabled us to address whether each of the 14 TFs are involved in regulating genes that are induced or repressed in response to ABA (Figure 2). Moreover, this analysis revealed that two known regulators of ABA signaling, ABF1 and ABF3 (Choi et al., 2000), are at the top of the list for having targets that are highly enriched for the ABA induced genes (Figure 2). Next, we further separated the TF-regulated targets of each of the 14 TFs into TF-induced or TF-repressed target sets using the *Query* function of ConnecTF. This analysis enabled us to determine the TF-target *specificity* (e.g. percent of TF-regulated targets that are ABA responsive), TF-target *influence* (e.g. percent of ABA responsive genes regulated by each TF), and *P*-value of the overlap of TF-target genes with induced and repressed ABA responsive genes (Supplemental Table 4). This analysis revealed that for the majority of the 14 TFs, the TF-induced targets overlap significantly with genes induced by the ABA signal, while TF-repressed targets overlap significantly with the genes repressed by ABA treatment.

**Figure 2.**
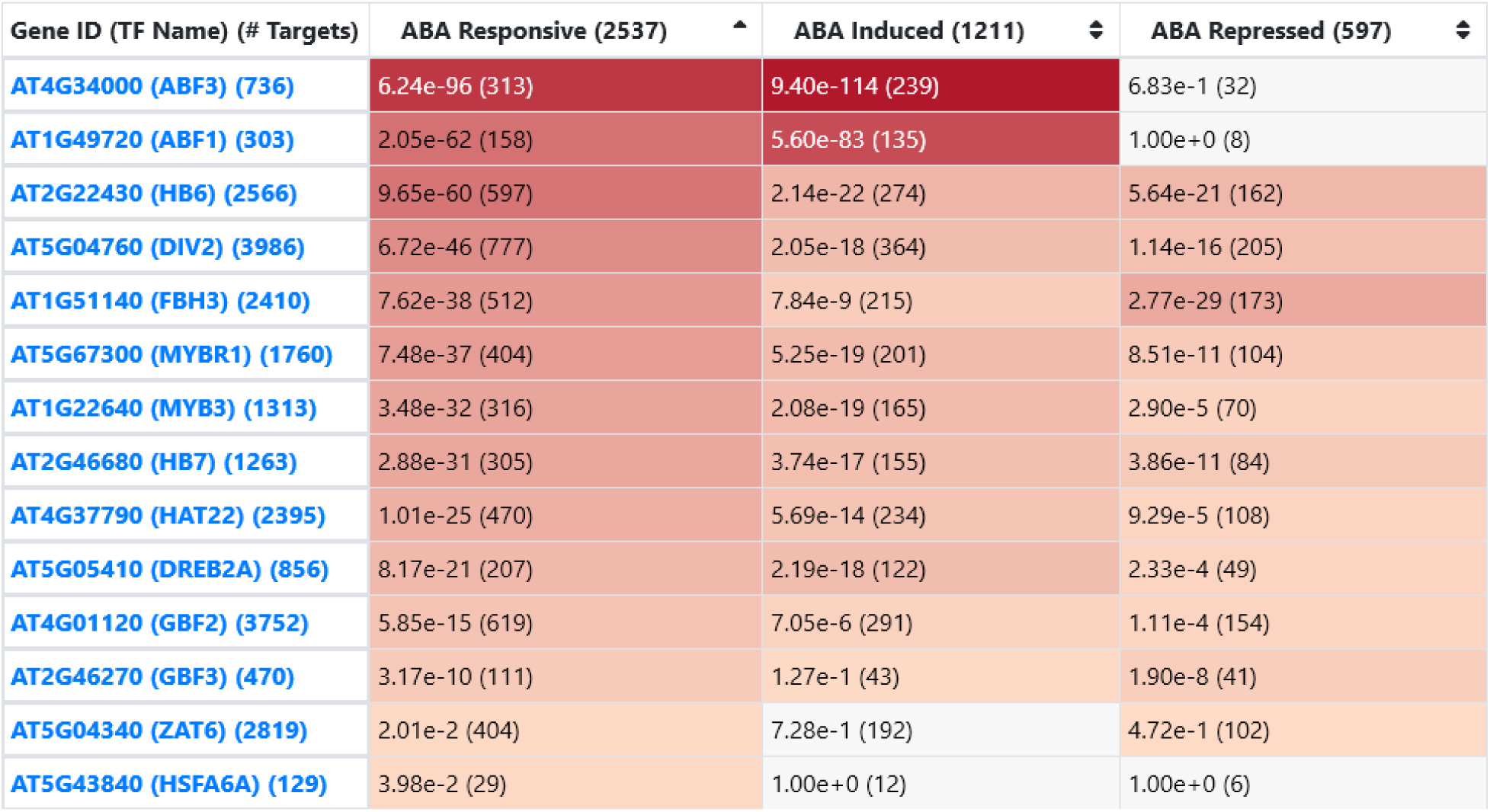
Case Study 1: Ranking significance of 14 TFs in regulation of ABA responsive genes. ConnecTF was used to address whether the direct regulated targets of 14 ABA responsive TFs identified in isolated root cells using the TARGET assay (Supplemental Table 3) are enriched for ABA responsive genes identified in Song et al. (Song et al., 2016). This screenshot from the ConnecTF website shows the results of the *Target List Enrichment* tool. We observed that the validated regulated targets of each of the 14 TFs are enriched for ABA responsive genes, including either ABA induced genes or ABA repressed genes (*P*-value < 0.05, Fisher’s exact test). Known ABA regulators ABF1 and ABF3 (Choi et al., 2000) are among the most enriched and are primarily involved in regulating targets that are induced in response to ABA treatment.

#### Distinct cis-motifs are enriched in the TF-induced vs. TF-repressed targets of 14 TFs in ABA signaling

We next sought to use the TF-target gene binding and TF-target gene regulation data for these 14 TFs to determine whether the TFs act alone, or in combination, to regulate the target genes in the ABA response pathway. To this end, we first asked whether the validated cis-binding motif for each TF (collected from Cis-BP) (Weirauch et al., 2014) showed specific enrichment in either the TF-induced or the TF-repressed target gene lists, as we found in a previous study of 33 TFs (Brooks et al., 2019). To do this, we first made a query in ConnecTF that returns the TF-induced or TF-repressed targets for each TF as separate gene lists. Next, we selected the *Individual Motifs* tab from within the *Motif Enrichment* results page. The default setting returns the cis-element enrichment in the 500 bp promoter region of the validated target genes of a TF for any cis-motif for that TF. Users can also define other genic regions of target genes (2000 bp promoter, 1000 bp promoter, 5’ untranslated region (UTR), coding sequence (CDS), introns, 3’ UTR and exons), or choose a cis-motif for another TF, e.g. a putative partner, and ConnecTF will calculate enrichment for the selected motif(s) in the selected genic region(s).

For the 14 TFs in the ABA pathway, we examined their TF-induced vs TF-repressed gene target lists for enrichment of their own cis-motif and show examples for the TFs HB7, MYB3 and ZAT6, (Figure 3, see Supplemental Data for query used to generate this figure). We found that a majority of the 14 TFs tested have enrichment of their known cis-element in *either* their induced or repressed targets that we identified as directly regulated TF-targets in root cells (Supplemental Table 5). Of these, 7/14 TFs (including HB7, Figure 3A) show enrichment of at least one known cis-motif for that TF exclusively in the TF-induced targets, while 2/14 (MYBR1 and MYB3, Figure 3B) show specific enrichment of cis-motif for that TF exclusively in the TF-repressed targets (Supplemental Table 5). For 5/14 TFs (including ZAT6, Figure 3C), there was no enrichment of their known cis-binding motif in either the TF-induced or TF-repressed targets.

**Figure 3.**
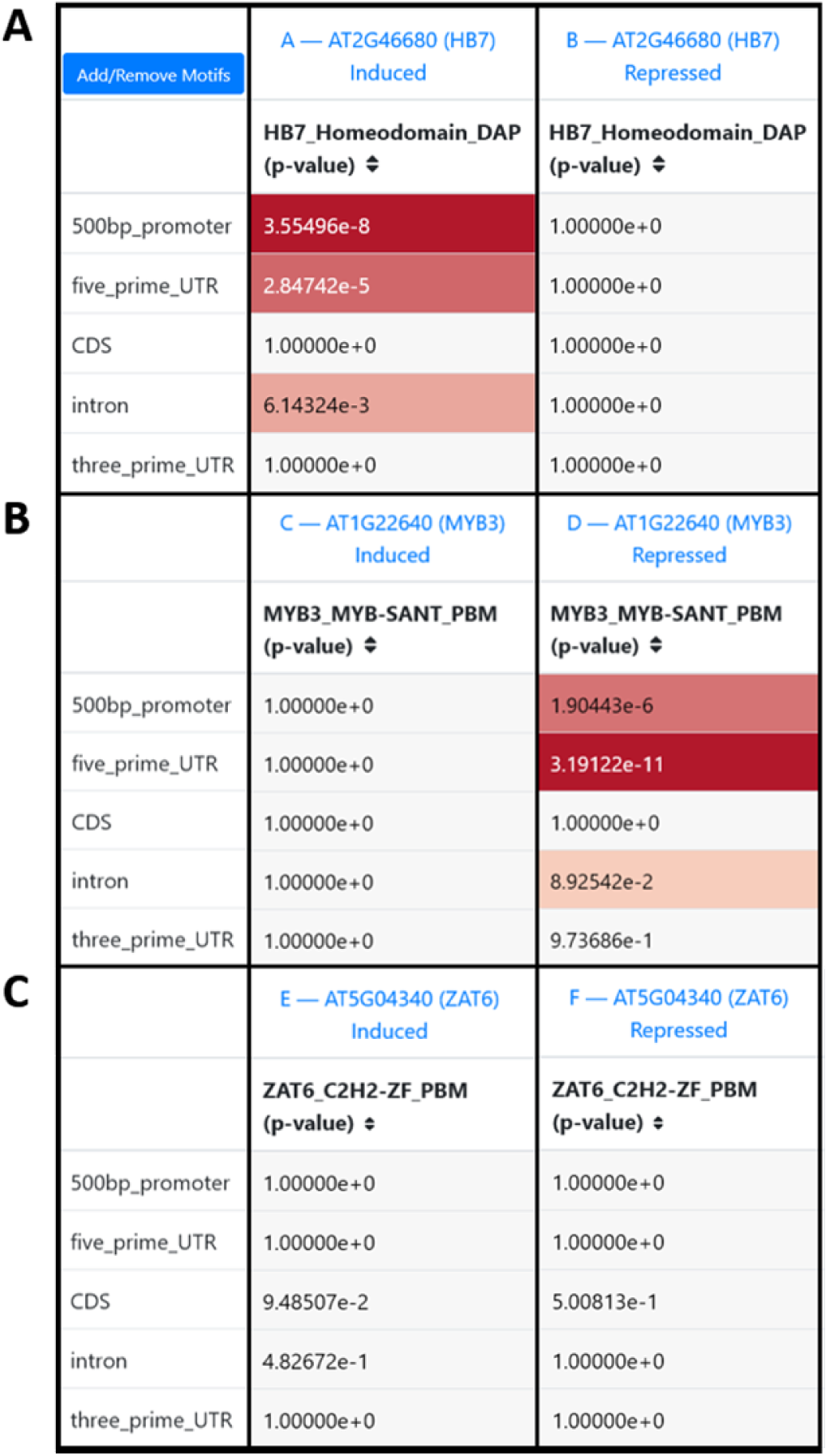
Case Study 1: Known cis-binding motifs for a TF are enriched in specific subsets of TF-regulated genes (induced vs. repressed). The ConnecTF database houses 1,310 experimentally determined cis-binding motifs for 730 Arabidopsis TFs and 37 cis-binding motifs for 26 maize TFs (Table 1 and Supplemental Table 1). Users can use this resource determine if any of these cis-motifs are enriched in the targets of the queried TF(s) using the *Individual Motifs* section of the *Motif Enrichment* tab. Here, we present a screenshot demonstrating how ConnecTF can be used to determine the enrichment of cis-motifs within the subset of targets of a TF (e.g. TF-induced or TF-repressed targets). The results show that the A) the HB7 cis-motif is enriched only in the TF-targets induced by HB7 in a root cell-based TF-assay, but not in the targets whose expression is repressed by HB7, B) the MYB3 cis-motif is enriched only in the TF-targets repressed by MYB3, but not the MYB3-induced targets, and C) the known motif for ZAT6 is not found to be enriched in either the induced or repressed targets of ZAT6. *P*-values were calculated using the Fisher’s exact test.

While cis-motif enrichment indicates where a TF is *likely* to directly bind in the genome, validated direct binding to specific genomic loci is available from in vitro TF-target gene binding (e.g. DAP-seq experiments) housed in the ConnecTF database (O’Malley et al., 2016). For the 5/14 ABA responsive TFs for which DAP-seq data is available (FBH3, GBF3, HB6, HB7, and MYBR1), our comparison of TF-induced or TF-repressed targets with in vitro TF-bound targets supported the cis-motif enrichment results. That is, for FBH3, HB7 and HB6, only the TF-induced target gene lists were enriched for genes that were bound in vitro to that TF, while for MYBR1, only TF-repressed targets were enriched in genes that were bound in vitro to that TF (Supplemental Table 6). GBF3, which had no cis-motif enrichment in either the TF-induced or TF-repressed directly regulated targets, also had no enrichment of TF-binding in vitro in either set of TF-regulated targets (Supplemental Table 6).

#### TF-regulated genes are largely TF-bound, while TF-bound genes are infrequently TF-regulated

An outstanding question related to TF-target validation datasets, is when and whether TF-binding results in gene regulation. To answer this question, we asked whether genes that are bound by each of the 14 ABA responsive TFs in planta, based on ChIP-seq experiments (Song et al., 2016), significantly overlap with either TF-induced or TF-repressed genes identified in root cells (Supplemental Table 3). To do this, we used the *Gene Set Enrichment* tool in ConnecTF, which reports whether the pairwise overlap between any two queried experimental analyses is greater or less than expected by chance (Fisher’s Exact test) (Figure 4D). This *Gene Set Enrichment* function is based on the Genesect tool in VirtualPlant (Katari et al., 2010) and described in Krouk et al. (Krouk et al., 2010). As an example, for three TFs - HB7, MYB3 and ZAT6 - the *Gene Set Enrichment* results show that both the TF-induced and TF-repressed target gene lists significantly overlap with the TF-bound targets of that TF (*P*-value<0.05, Fisher’s exact test) (Figure 4, see Supplemental Data for query used to generate this figure). Extending this analysis to all 14 ABA responsive TFs, we find that 9/14 TFs have a significant overlap of TF-bound genes in planta with both the list of TF-induced and TF-repressed targets of that TF, as validated in root cells (*P*-value<0.05, Fisher’s exact test) (Supplemental Table 7). For 4/14 of the TFs - ABF1, ABF3, DREB2A and HSFA6A - we found a significant overlap of the TF-bound targets only with the TF-induced targets (*P*-value<0.05, Fisher’s exact test). By contrast, only 1/16 TFs (GBF2) had a significant overlap of TF-bound targets only with the list of TF-regulated targets that are repressed (*P*-value<0.05, Fisher’s exact test) (Supplemental Table 7).

**Figure 4.**
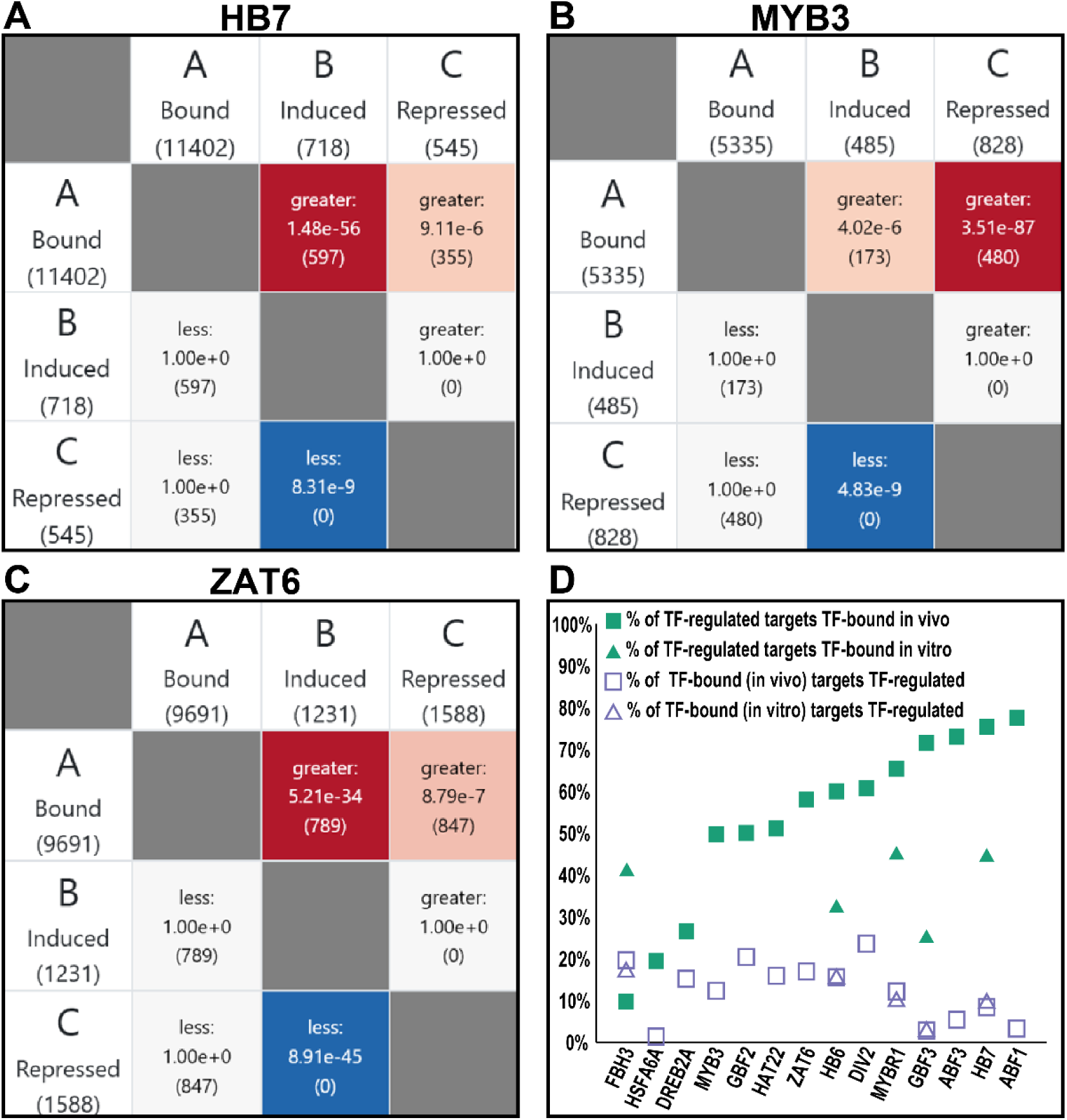
Case Study 1: TF-regulated gene targets are largely TF-bound, while TF-bound genes are infrequently TF-regulated. The *Gene Set Enrichment* tool in ConnecTF can be used to determine if the pairwise overlap of the target gene lists of two TF analyses is significant (Fisher’s exact test). This feature enables users to answer common questions such as “What is the overlap between ChIP and TF perturbation of the same TF? Or, how significant is the overlap of the targets of two different TFs?” To demonstrate this feature, for A) HB7, B) MYB3 and C) ZAT6, we show screenshots from the ConnecTF site of the overlap between bound targets as determined by in planta ChIP (Song et al., 2016) and the induced and repressed TF-targets that we determined in isolated root cells in this study using the TARGET assay. For each TF, the bound targets significantly overlap with both the TF-induced and TF-repressed targets identified in cells. D) When we performed this overlap of TF-regulation and TF-binding for all 14 TFs (Supplemental Tables 6 and 7), we observed that the percent of TF-regulated genes that are TF bound is much greater than the percent of TF-bound genes that are TF-regulated, regardless of whether the binding data is in vivo or in vitro. This suggests that TF-biding is a poor indicator of gene regulation in the absence of complimentary TF-regulation data for each TF.

Finally, we used ConnecTF to evaluate the relationship of TF-binding vs. TF-regulation datasets. Overall, our integrated analysis of TF-binding and TF-regulation data showed that for 11/14 of the ABA responsive TFs, greater than 50%, and as much as 75%, of TF-target genes that were TF-regulated in root cells were also bound by that TF in planta (Figure 4D). By contrast, for all 14 TFs, the number of TF-bound targets in planta that were regulated by that TF never exceeded 25% (Figure 4D).

#### Identifying partner TF_2_-binding motifs in TF_1_-regulated genes

Next, for each set of TF_1_-regulated targets (either induced or repressed) that showed no enrichment of the known cis-binding motif for TF_1_ (Supplemental Table 5), we used ConnecTF to search for overrepresentation of cis-motifs for potential partner TF_2_s in those sets of TF_1_-regulated genes. Rather than searching for all 1,310 cis-motifs available for Arabidopsis from CIS-BP (Weirauch et al., 2014), we limited our search to the 80 cis-motif clusters generated from all available Arabidopsis thaliana cis-motifs (Brooks et al., 2019), now housed in the ConnectTF database.

First, we performed cis-motif enrichment analysis on the validated target gene lists of three TFs - HB7, MYB3 and ZAT6 (Figure 5). For each of these TFs, we hypothesized that they could act directly on gene targets, or through a TF_2_ partners, based on our analysis of TF-regulation, TF-binding and cis-motif enrichment. For HB7, while both HB7-induced and HB7-repressed targets identified in root cells are each bound by HB7 in planta (by ChIP-seq) (Figure 4A), the known HB7 cis-motif is only enriched in the HB7-induced targets (Figure 3A). Using ConnecTF cis-analysis functions, we found that the HB7-repressed target gene list is enriched in a cis-motif (cis-cluster 13) for a WRKY TFs (*P*-value<0.05, Fisher’s exact test) (Figure 5A). This finding suggests HB7-repression of gene targets is mediated by one or more TF_2_ partners in the WRKY TF family. For MYB3, while both MYB3 induced and repressed target gene lists identified in root cells are each enriched in genes bound by MYB3 in planta (e.g. ChIP-seq) (Figure 4B), the MYB3 cis-motif is only enriched in the list of MYB3-repressed targets (Figure 3B). By contrast, the list of MYB3-induced targets are enriched in cis-motifs (cis-clusters 6, 39, 68) for TF_2_s in the bZIP/bHLH/BZR and CAMTA/FAR1 TF families (*P*-value<0.05, Fisher’s exact test) (Figure 5B). This result suggests that MYB3 induces target genes via an indirect interaction with TF_2_(s) from the bZIP1/bHLH/BZR, or CAMTA/FAR1 families. Lastly, although the list of ZAT6-induced and ZAT6-repressed targets in root cells are enriched in genes bound by ZAT6 in planta (e.g. ChIP-seq) (Figure 4C), there is no enrichment of the known ZAT6 cis-element in either set of ZAT6-regulated genes (Figure 3C). Instead, the list of ZAT6 induced genes are enriched is cis-elements for cis-clusters 6 and 39 from the bZIP/bHLH/BZR TF families, while the list of ZAT6-repressed genes are enriched in cis-cluster 13 for WRKY TFs (*P*-value<0.05, Fisher’s exact test) (Figure 5C). This suggests that ZAT6 regulates both its induced and repressed targets via TF_2_(s) in these families.

**Figure 5.**
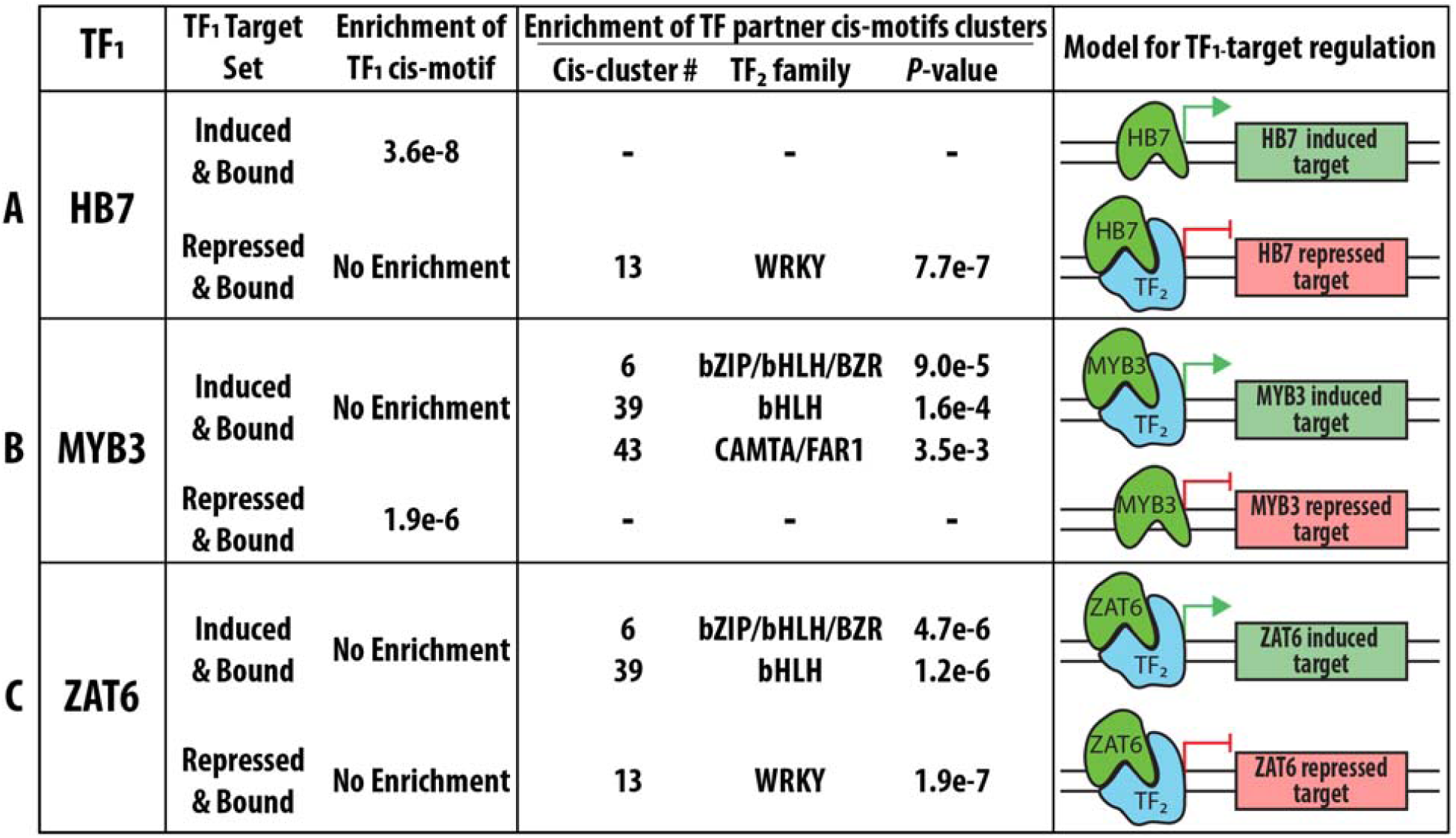
Case Study 1: Putative cis-motifs for TF_2_ partners are identified in indirectly bound TF_1_-targets. ConnecTF was used to combine the new TF-regulation data generated in this study for 14 ABA responsive TFs with existing TF-binding data in planta (Song et al., 2016), and to reveal mode-of-action for how these TFs function to regulate target genes in the ABA signaling pathway. Here we summarize these results for 3/14 TFs; A) HB7, B) MYB3, and C) ZAT6. For both HB7 and ZAT6, we found that TF-repressed and TF-bound targets, which lack enrichment of the known cis-motif for that TF (see Figure 3), had enrichment of the cis-motif cluster representing WRKY TFs (Brooks et al., 2019). Similarly, for MYB3 and ZAT6, the TF-induced and TF-bound targets that were not enriched in the cis-motif for these TFs, were each enriched for cis-motif clusters 6 and 39 which represents the bZIP/bHLH/BZR families of TFs (Brooks et al., 2019). This cis-analysis allowed us to derive a model for each TF (e.g. HB7, MYB3 and ZAT6) which describes how physical interactions with putative partner TFs (TF_2_s) enable the TF to regulate subsets of its target genes, even in the absence of direct binding.

When we analyzed all 14 TFs using this approach, we observed that cis-motif clusters 6 and 39 are enriched (*P*-value<0.05, Fisher’s exact test) in the lists of TF-induced or TF-bound gene targets of 7/14 of the ABA-responsive TFs (Supplemental Table 8). Furthermore, we found that cis-motif Clusters 6 and 39 are enriched in the list of genes induced by ABA (*P*-value<0.05, Fisher’s exact test), but not in the list of ABA-repressed genes (Supplemental Table 8). This result suggests that partner TF_2_s from the bHLH/bZIP/BZR TF family/families work with MYB3, ZAT6, and other ABA-responsive TFs to regulate these ABA-responsive targets. Likewise, cis-motif cluster 13 which represents WRKY TFs, is enriched in the list of the TF-repressed or TF-bound targets of 7/14 TFs, as well as in the list of genes that are repressed in response to ABA (*P*-value<0.05, Fisher’s exact test) (Supplemental Table 8). Thus, these studies uncovered potential TF_2_ partners of 14 TFs involved in the ABA response.

### Case Study 2: Refining/pruning inferred gene regulatory networks using validated TF-target data

In this case study, we show how ConnecTF can be used to readily evaluate the relevance of, and combine gold-standard TF-target gene validation data for refining network predictions using automated precison/recall analysis. This feature will advance the systems biology cycle of network prediction, validation, and refinement.

#### Automated precision/recall analysis and refinement of a nitrogen response GRN

As an example, we show how ConnecTF can automate a precision/recall analysis on a GRN inferred from time-series transcriptome data of the nitrogen response in Arabidopsis roots (Brooks et al., 2019). As a gold standard validation data, we selected the TF-target regulation data based on TF-perturbation experiments in root cells using the TARGET system (Bargmann et al., 2013). This set of 55 TFs includes the 33 nitrogen response TFs from Brooks et al. (Brooks et al., 2019), 8 TFs from Alvarez et al. (Alvarez et al., 2020), and the 14 ABA response TF-target regulation datasets generated in root cells in this study (Supplemental Table 3). To initiate this precision/recall analysis of the inferred nitrogen response GRN in ConnecTF, we first queried the 55 TF-target gene regulation datasets performed in root cells using the *Query* page. To determine which of these 55 TFs were relevant to our GRN analysis, we used the *Target Network* box to select the “Root Predicted Nitrogen Network” from Brooks et al. (Brooks et al., 2019). This query returned a total of 32 TFs and 1,349 validated TF-target genes in the predicted nitrogen-regulatory network. This query automatically generates a precision/recall curve, which is seen in the AUPR section at the bottom half of the *Network* tab (Figure 6, see Supplemental Data for query used to generate this figure). The slider or textbox above the AUPR plot can be used to select a precision cutoff score, which will update the interactive AUPR graph and table with details of a pruned/refined network, e.g. the predicted TF-target edges whose score equals or exceeds the selected precision score threshold. In this example, the selected cutoff of 0.32 reduced the size of the predicted N-regulatory GRN from 240,410 interactions between 145 TFs and 1,658 targets to a refined high-confidence GRN of 4,343 interactions between 143 TFs and 215 target genes whose predicted interactions passed the threshold set by the precision/recall analysis of the validated TF-target gene interactions.

**Figure 6.**
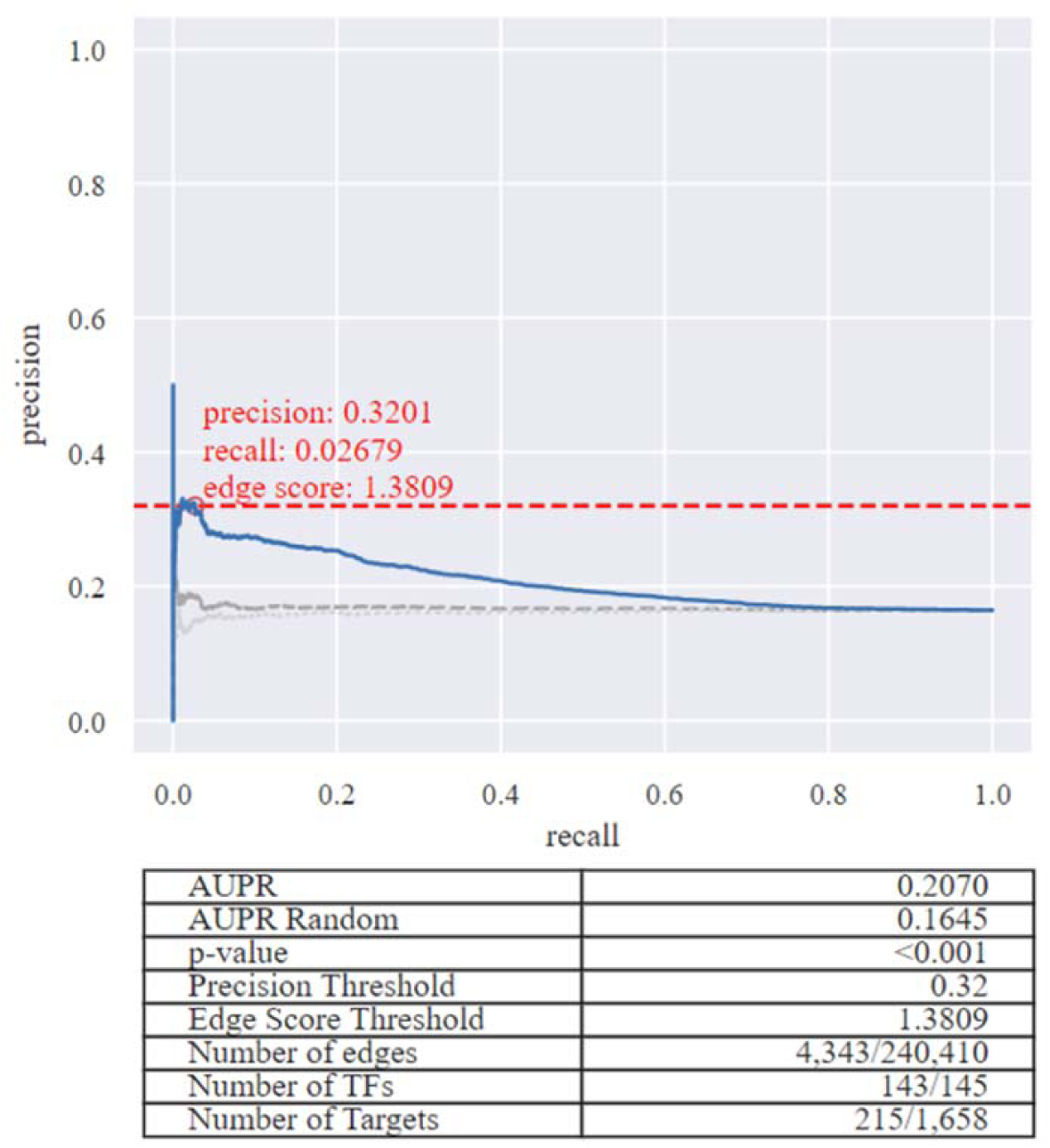
Case Study 2: Performing an automated precision/recall analysis on an inferred network uploaded by the user. Users are able to perform an automated precision/recall analysis on a predicted network. To do this, the user first uploads a ranked list of TF-target interactions in a predicted networks into ConnecTF from the *Query* page using the *Target Network* box. Next, they can validate the predicted network using TF-target gene validated data in the ConnecTF database. Once they do this, within the *Network* tab, a precision/recall analysis (AUPR) section will be automatically generated for the predicted network, using selected TF-target validation datasets in the ConnecTF database, and display a precision/recall plot and summary table. The user can then select a precision cutoff using the sliding bar above the plot, which will interactively update the AUPR graph, summary table, and the network that is visualized or exported. Query filters enable the user to select which TFs and the specific types of edges that should be used as the “gold standard” to perform precision/recall analysis of the predicted network. Here we show a screenshot for an example where we used the time-based inferred network from Arabidopsis roots (Brooks et al., 2019), and all validated edges from TFs whose TF-regulated targets were identified in root cells (39 experiments) to demonstrate this AUPR feature of ConnecTF.

GRNs constructed based on co-expression data can also be validated in a similar manner. To this end, we provide a precision/recall example for a GRN built from the co-expression network available in the Atted-II database (Obayashi et al., 2018). We pruned this co-expression GRN using all TF-regulation data in the ConnecTF database (Supplemental Figure 2, see Supplemental Data for query used to generate this figure).

Using the appropriate buttons at the top of the *Network* page, the user can download the pruned/refined network as a network file (in JSON or SIF formats) or visualize the network in the browser (*Open Network*). The precision cutoff can be further modified while viewing the network in the browser using the slider or text box in the *Additional Edges* menu. Edges within the network can be hidden to highlight a specific interaction type of interest (e.g. time-based edge predictions) or additional edges can be added from a file the user uploads. The resulting pruned network can be saved as a JSON file or an image exported.

#### TF-regulation data outperforms in vitro TF-binding as a gold-standard for precision/recall analysis

Next, we demonstrate how ConnecTF can be used to evaluate which TF-target validation datasets are most effective for use as gold standards for network refinement. As an example, the automated functions in ConnecTF enabled us to rapidly evaluate and compare the relative AUPR performance of different TF-target validated datasets (e.g. TF-binding (DAP-seq) vs. TF-regulation) in precision/recall analysis of a GRN inferred from time-series nitrogen response in Arabidopsis roots (Brooks et al., 2019). The TF-target validated datasets we tested are; 1) TF-Regulated gene sets: TF-target sets regulated in root cells (e.g. TARGET assay) (Brooks et al., 2019; Alvarez et al., 2020), 2) TF-Bound gene sets: TF-target sets bound in vitro (DAP-seq) (O’Malley et al., 2016), or 3) TF-regulated and TF-bound gene sets: TF-target sets regulated in root cells (TARGET assay) and bound in vitro (DAP-seq) (Table 2). For the gene sets that involved TF-target binding (i.e. 2 and 3 above), we also used the DHS data (Sullivan et al., 2014) housed in the ConnecTF database to filter for DAP-seq peaks that occur in open chromatin regions in root tissue.

**Table 2.** Precision/recall analysis of a GRN inferred network from time-series nitrogen response data in Arabidopsis roots (Brooks et al., 2019) performed using automated precision/recall functions in ConnecTF using different sets of experimentally validated edges in the ConnecTF database.

| **Validated Edges Used** | **AUPR** | **AUPR randomized network** | **p-value** | **Percent improvement vs. random** |
| --- | --- | --- | --- | --- |
| TF-Regulated only (TARGET) | 0.2025 | 0.1595 | <0.001 | 27% |
| TF- bound only (in vitro) (DAP-seq) | 0.3257 | 0.2967 | <0.001 | 10% |
| TF-Regulated and TF- bound (in vitro) (TARGET ∩ DAP-seq) | 0.0863 | 0.0614 | <0.001 | 41% |
| TF-bound only (in vitro) (DAP-seq)/ DHS filtered (root) | 0.1908 | 0.1682 | <0.001 | 13% |
| TF-Regulated and TF-bound (in vitro)/ DHS filtered (root) (TARGET ∩ DAP-seq) | 0.0555 | 0.0398 | <0.001 | 39% |

By comparing the precision/recall results on networks refined using these three validated TF-target gene datasets, we found that using TF-regulated target data identified in root cells as “gold standard” resulted in a higher AUPR, and greater improvement in AUPR relative to the randomized predicted network, compared to using in vitro TF-binding target data alone (DAP-Seq) (Table 2). Also, we found that combining TF-target regulated and TF-target bound datasets reduced the AUPR, however, it resulted in a greater improvement relative to the randomized network, compared to using TF-regulation datasets only. Finally, we found that applying the DHS filter to DAP-seq peaks reduced the AUPR, and had only a small effect on the improvement of the AUPR relative to the randomized network, compared to the same set of edges without the DHS filter (Table 2). Thus, the ability to test and combine TF-target datasets in an automated AUPR analysis enabled us to rapidly determine which datasets were most effective for use in network refinement.

### Case Study 3: Charting a network path by combining validated TF-target data for multiple TFs

An important feature that distinguishes ConnecTF from most other available analysis tools/platforms concerning TFs, is its *Query* building function. The *Query* builder allows users to readily select, parse, and combine TF-target gene validation data from different TF experiments and research groups. For example, below we demonstrate how ConnecTF can be used to chart a network path from the direct targets of a TF_1_ to its indirect targets via secondary TFs (TF_2_s). We initially conceived of this Network Walking approach which we manually executed in Brooks et al. (Brooks et al., 2019). As an example, we show how ConnecTF can be used to chart a network path from TF_1_ - NLP7, a master TF in the nitrogen signaling pathway – to its direct TF_1_-targets to its indirect targets, by combining TF-target regulation and TF-target binding datasets from two different NLP7 studies (Marchive et al., 2013; Alvarez et al., 2020).

#### Step 1. Identify direct vs. indirect targets of TF_1_

The first step in charting a network path is to identify the direct vs. indirect targets of TF_1_. To this end, we used the *Query* function in ConnecTF to identify direct NLP7 (TF_1_) targets as genes that are both NLP7-regulated and NLP7-bound (Marchive et al., 2013; Alvarez et al., 2020). Next, we identified indirect NLP7 targets as genes that are regulated, but not bound by NLP7 in ChIP experiments (Marchive et al., 2013; Alvarez et al., 2020). We executed two simple queries in ConnecTF to produce these lists of direct targets of NLP7 (Figure 7A, Query 1, see Supplemental Data for details of Query 1) and indirect targets of NLP7 (Figure 7A, Query 2, see Supplemental Data for details of Query 2). The list of genes resulting from these queries can be saved within ConnecTF, to be used as direct vs. indirect target gene lists of the TF_1_ (NLP7) for further analyses in the following steps, or downloaded by the user.

**Figure 7.**
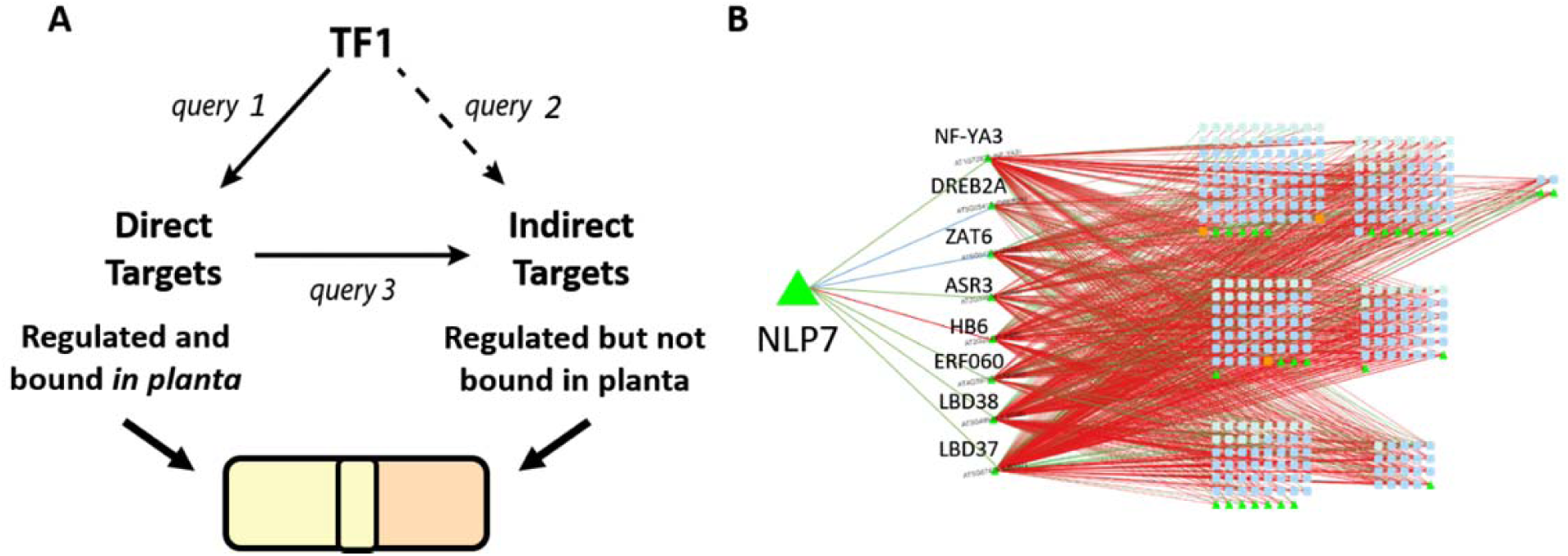
Case Study 3: Network Walking: Using ConnecTF to chart a network path from TF_1_ → TF2s → indirect targets of TF_1_. The query system of ConnecTF can be used in an iterative process, with the results of one query being used to filter the TFs and/or target genes of other queries. This facilitates the building of more complex GRNs, such as charting a network path from TF_1_ to its downstream TF_2_s and indirect targets. A) ConnecTF can be used to chart a network path from a TF_1_ via its direct TF_2_s to its indirect targets using the Network Walking approach described in Brooks et al. (Brooks et al 2019). Simple queries can be used in ConnecTF to integrate TF-target binding and TF-target regulation datasets to identify TF_1_ direct targets (TF_1_-regulated and TF_1_-bound, query 1) and TF_1_ indirect targets (TF_1_-regulated but not TF_1_-bound, query 2). The results of a query can also be saved and used to filter subsequent user queries, as in query 3. B) We demonstrate the process of Network walking using NLP7, a master TF_1_ involved in nitrogen signaling, identifying a set of 8 direct intermediate TF_2_s targets acting downstream of NLP7 that control 68% of the NLP7 indirect targets.

#### Step 2. Connect TF_1_ to its indirect targets via its direct intermediate TF_2_s

With the lists of direct vs. indirect targets of a TF_1_ (NLP7), we can now perform the second step of charting a network path in the Network Walking approach. In Step 2, we used ConnecTF to connect the indirect targets of NLP7 via TF_2_s that are themselves direct targets of NLP7. To do this, we queried all the TF-target regulation datasets performed in root cells (55 TFs) in the ConnecTF database, restricting the results returned to the indirect targets of TF_1_ (e.g. NLP7 regulated, but not bound) using the *Target Genes* filter on the query page. For this query, we also restricted the TF_2_s to the direct targets of NLP7, as identified in Step 1, using the *Filter TFs* option (Figure 7A, Query 3, see Supplemental Data for details of Query 3). The resulting *Table* tab shows the complete set of validated TF-target edges from 8 TF_2s_ that are direct targets of NLP7 (e.g. TF_2_s: ASR3, NF-YA3, DREB2A, ZAT6, ERF060, HB6, LBD37 and LBD38) to NLP7 indirect targets. From the *Target Enrichment* tab, we see that all 8 TF_2_s are enriched for NLP7 indirect targets (*P*-value<0.05, Fisher’s exact test), with NF-YA3, LBD37 and LBD38 being the most important based on TF-influence, target specificity and *P*-value of the overlap (Supplemental Figure 3, see Supplemental Data for query used to generate this figure).

#### Step 3. Visualizing the Network Path from TF_1_ → direct TF_2_(s) → indirect targets of TF_1_

Finally, we can visualize the resulting Network path from TF_1_ (NLP7) → 8 direct TF_2_ targets → indirect TF_1_ targets. We can do this in ConnecTF by going to the *Network* tab and clicking *Open Network* which will launch Cytoscape.js (Franz et al., 2015). Basic Cytoscape functionality is available within ConnecTF for viewing and adding additional edges to the network (Figure 7B), or the network can be downloaded as a JSON file and further modified by the user.

## Discussion

As the cost of Next-generation sequencing technologies declines and new methods are developed to identify/validate TF targets, computational tools to integrate the increasing amount and types of experimental data that relate TFs with their target genes are becoming increasingly important (Grossman, 2019). Enabling researchers not only to access these various types of TF-target validation datasets, but to perform analyses that integrates multiple datasets and multiple types of data, will further our understanding of the mechanisms by which TFs function alone and together in a GRN that affects a biological pathways of interest.

To this end, we developed ConnecTF (https://ConnecTF.org) to facilitate these types of research questions in GRN analysis/validation. Moreover, we have designed ConnecTF to be accessible to biologists with a wide-range of computational skills. As a resource for the plant research community, we are hosting two versions of ConnecTF for Arabidopsis and maize, with a combined 4,577,488 edges for 562 TFs (Table 1 and Supplemental Table 1). In our three case studies, we provide examples of how ConnecTF can enable an integrated analysis of TF-target gene interactions that lead to biological insights of TF modes-of-action, using GRNs involved in the ABA and nitrogen response pathways.

The ConnecTF database was designed to specifically house large-scale datasets for TF-binding and TF-regulation. For Arabidopsis, the vast majority of data for TF-target binding is in vitro (387 TFs) (O’Malley et al., 2016), and a more limited set of large-scale TF-target binding datasets in vivo (26 TFs) (See Table 1). The ConnecTF database houses the largest set of TF-target regulation data based on a high throughput TF-assay performed in isolated plant cells (58 TFs) (Varala et al., 2018; Brooks et al., 2019; Alvarez et al., 2020), which also includes new data on TF-target regulation for 14 TFs identified in this study (Table 1). The ConnecTF database also houses cis-motif data for 730 Arabidopsis TFs (Weirauch et al., 2014). Finally, the database contains information on TF-TF protein interactions (Yazaki et al., 2016; Trigg et al., 2017), and the ability for users to filter in vitro TF-binding data for peaks occurring in open chromatin regions from different tissues identified using ATAC-seq (Lu et al., 2019) or DHS (Sullivan et al., 2014).

In Case Study 1, we used ConnecTF to combine TF-target gene validation data for 14 TFs in the ABA signaling pathway for which we have datasets for TF-binding in vivo (14/14 TFs) (Song et al., 2016), TF-regulation in root cells (14/14 TFs) (Supplemental Table 3), TF-binding in vitro (DAP seq) (5/14 TFs) (O’Malley et al., 2016), and cis-motif data (14/14 TFs) (Weirauch et al., 2014). Our integrated analysis of this TF-regulation and TF-binding data using ConnecTF allowed us to discover that TF-regulation is a good indicator of TF-binding, but TF-binding is a poor indicator of TF-regulation. Specifically, up to 78% of the direct TF-regulated genes were TF-bound in planta (Figure 4D and Supplemental Table 7). However, the reverse is not the case, as for these 14 TFs, at most 24% of TF-targets bound in planta were TF-regulated in root cells (Figure 4D and Supplemental Table 7). While this could be due to the different systems used in this study, TF-binding is known to be a poor indicator of TF-regulation across many organisms, even when TF regulation and TF binding are compared from the same tissue (Phuc Le et al., 2005; Bolduc et al., 2012; Arenhart et al., 2014), or even the same cell samples (Para et al., 2014).

Using ConnecTF to readily intersect TF-bound and TF-regulated gene targets for a large number of TFs also allowed us to develop mode-of-action models for how TF binding might lead to induction or repression of target genes by the TF. For this analysis, we use the direct regulated TF-targets validated in a plant cell-based system comprising 58 TFs, including 14 TFs from this current study (Supplemental Table 3) and 44 TFs from our previous work (Varala et al., 2018; Brooks et al., 2019; Alvarez et al., 2020). Combined, these TF-target regulation datasets have shown that 57/58 TFs can act as both an inducer and repressor, depending on the target genes. The one exception is HSFA6A, which acted primarily as an inducer (127 genes), and down-regulated only two targets (Supplemental Table 3). We also observed that the known cis-binding motif for a TF is most often significantly enriched in either the induced or repressed targets of that TF (Figure 3 and Supplemental Table 5), as we saw previously for 11 TFs (Brooks et al., 2019). This broader finding indicates that direct binding of a TF to its targets most often has a specific effect on target gene expression (e.g. either induction or repression, depending on the TF). Importantly, our integrated data analysis showed that TF-TF interactions likely play a role in the “switch” of a TF from an activator or repressor, depending on the target gene (Figure 5), as described below.

The simplest model for TF-target regulation is through direct interaction of a TF via DNA-binding to cis-regulatory regions in its target genes. However, it has been observed that cellular and genomic context, including TF-TF cooperativity, can play an essential role in how a TF controls target gene expression (Yáñez-Cuna et al., 2012; Para et al., 2014; Slattery et al., 2014; Alvarez et al., 2020; de Boer et al., 2020). Indeed, we found examples of regulation of TF-target gene expression in the absence of evidence for direct TF-binding (Figures 3-5, Supplemental Tables 5-7). In these cases, TF regulation of the target gene could occur by indirect TF_1_ binding to a target via its association with partner TF_2_s, sometimes referred to as “tethering” (Stender et al., 2010). Previous studies have compared ChIP and DHS foot-printing to distinguish between direct and indirect TF-target binding (Gordân et al., 2009; Neph et al., 2012). In case study 1, we demonstrate that using ConnecTF to integrate TF-target binding and TF-target regulation data enabled us to discover that for a majority of the 14 TFs in the ABA signaling pathway, both their TF-induced and TF-repressed target gene sets overlap significantly with in planta bound targets (Figure 4 and Supplemental Table 7). This occurs even when evidence for direct TF-binding, in the form of cis-motif enrichment or in vitro TF-binding, is absent (Supplemental Tables 5 and 6). Moreover, we used ConnecTF to identify potential partner TF_2_s involved in the indirect target binding of TF_1_, by enrichment of cis-binding motif clusters for other TF families (Brooks et al., 2019) in the direct regulated targets of the TF_1_s (Figure 5).

To do this, we looked for cis-motifs enriched in sets of TF_1_-regulated targets that are likely indirectly bound, i.e. those that are TF_1_-regulated (induced or repressed) and lack enrichment of the cis-motif for that TF_1_, but are bound to the TF_1_ in planta (e.g. by ChIP-seq) (Figure 5). For HB7, we found evidence for direct TF-binding leading to activation, while indirect binding leads to repression of its targets (Figure 5A). For MYB3, direct binding leads to target gene repression, while indirect binding leads to activation (Figure 5B). For ZAT6, we only found evidence for indirect binding to either its induced targets or repressed targets (Figure 5C). For the induced targets of ZAT6 (Figure 5C), we identified the enrichment of cis-motif clusters containing a core G-box motif, known to be bound by bZIP and bHLH TFs (de Vetten and Ferl, 1994; Toledo-Ortiz et al., 2003). Validation of this predicted TF-TF interaction leading to induction of ZAT6 indirect targets via a TF_1_-TF_2_ interaction comes from a known ZAT6 interaction with UPB1, an ABA-responsive bHLH TF (Trigg et al., 2017). This finding, uncovered using ConnecTF, suggests a simple explanation for how ZAT6 may induce target genes indirectly via a TF_1_-TF_2_ interaction (e.g. ZAT6-UPB1 complexes) (Figure 5). By contrast, cis-element analysis of the repressed targets of ZAT6 and other TFs (e.g. HB7) reveals an enrichment of cis-motif cluster 13 (Figure 5 and Supplemental Table 8), a core W-box motif known to be bound by WRKY TFs (Rushton et al., 1995).

A remaining question is how do 3/14 TFs tested in the ABA signaling pathway (ABF1, ABF3, and DREB2A) regulate target gene transcription without binding to those targets either directly or indirectly (Supplemental Tables 5 and 7)? These three TFs show no cis-enrichment in their repressed regulated targets of their own known cis-motif(s), nor do their TF-repressed target genes show enrichment of TF-bound targets in planta (Supplemental Tables 5 and 7). Other regulatory mechanisms for transcriptional control have been reported that do not involve TF binding, either direct or indirect, to target genes. This includes the destabilizing of transcriptional complexes by a TF, as seen for SPL9 repression of anthocyanin biosynthesis (Gou et al., 2011), and TFs sequestering components of a transcriptional activating complex (Nemie-Feyissa et al., 2014). While it is not possible to directly determine whether these types of mechanisms apply to the 14 TFs in ABA signaling used in our analysis, the results demonstrate how ConnecTF can be used to generate testable hypotheses by integrating TF-regulation and binding datasets.

In case study 2, we demonstrate how ConnecTF can be used to readily compare a predicted GRN against a set of validated TF-target gold standard interactions. With the large amount of TF-target validation data being generated, many sophisticated methods are being developed that use machine learning to predict GRNs from TF-target regulation and binding datasets (Marbach et al., 2012; Banf and Rhee, 2017; Mochida et al., 2018). However, a major bottleneck to this effort is the limited availability of validated TF-target edges, along with a clear understanding of what types of experimentally validated TF-target interactions are most useful to use as a gold standard for benchmarking inferred networks (Marbach et al., 2012; Banf and Rhee, 2017). The automated precision/recall analysis features of ConnecTF will contribute to overcoming this systems biology bottle neck by providing a resource for users to readily select and rapidly test which gold standard validated TF-target interactions are most useful to refine/prune or train their predicted networks.

We demonstrate how ConnecTF allows users to readily subset their gold standard validated TF-target edges based on specific criteria (e.g. edge type, *P*-value, fold change etc.), and compare how each subset affects the precision/recall analyses used to prune/refine inferred networks using automated functions in ConnecTF. We have incorporated features into ConnecTF that facilitate this functionality, including the ability to perform automated precision/recall analysis of user-provided ranked lists of inferred TF-target interactions in GRNs (Figure 6). In case study 2, we used these automated precision/recall analysis features to determine that TF-target regulation datasets are superior as gold-standard data, compared to in vitro TF-binding datasets. Specifically, we show that TF-target regulation datasets generated for 55 TFs using the TARGET cell-based TF-perturbation system (Bargmann et al., 2013; Varala et al., 2018; Brooks et al., 2019; Alvarez et al., 2020), results in a higher AUPR and statistical improvement relative to randomized networks, compared to using TF-target binding data from in vitro assays (i.e. DAP-seq), even when the same set of TFs are used (Table 2). Furthermore, the inferred network pruned with TF-targets that were *both* TF-regulated and TF-bound also resulted in a lower AUPR, compared to using all regulated targets for those TFs (Table 2). These results are unsurprising given what we observed in case study 1, that is, in vitro binding is extensive in the genome, but often represents only a subset of TF-regulated targets (Figure 4D). This is likely related to the observation that a majority of TF-binding in the genome does not result in gene regulation (Supplemental Table 6), and/or TF-TF interactions (i.e. indirect binding) which are not captured in this in vitro DNA binding assay.

In case study 3, we show how the ConnecTF platform enables users to integrate validated TF-target interactions from multiple TF datasets into a unified network path within a GRN, facilitating systems biology studies. To demonstrate this, we used ConnecTF to chart a network path that defined how NLP7, a master regulator of nitrogen signaling (Marchive et al., 2013; Alvarez et al., 2020), controls downstream genes through intermediate TF_2_s, following the Network Walking approach (Brooks et al., 2019). To do this, we showed how simple queries in ConnecTF can identify specific sets of targets of a TF_1_ (i.e. direct vs. indirect targets of TF_1_) and how these results can be combined with TF_1_ direct TF_2_ targets in an iterative process to chart network paths from TF_1_ (NLP7) → direct TF_2_s → direct targets of TF_2_s, which include indirect targets of TF_1_. Using ConnecTF allowed us to identify eight direct TF_2_ targets of NLP7 that are able to directly regulate 68% of NLP7 indirect targets (Figure 7). This network path shows that LBD37 and LBD39, which are known to be important in nitrogen uptake and assimilation in planta (Rubin et al., 2009), are the TF_2_s that are most influential on NLP7 indirect targets (Supplemental Figure 3). Thus, ConnecTF offers a way for users to identify the sequential action of TFs in a network path to regulate a pathway or set of genes of interest.

These three case studies are just some examples of the many ways that ConnecTF will be able to facilitate genomics and systems biology research in the plant community. We will host and maintain databases for the plant species Arabidopsis and maize. However, as we built the ConnecTF framework with common software packages and a species-independent structure, it is possible for users to easily set up an instance for any species of interest, and/or add new features and analysis tools. We provide detailed instructions on how to build private and/or public versions of ConnecTF for users interested in creating a database with their own data, and encourage other researchers to do so. As more TF-centric data is generated, we expect ConnecTF to be a powerful and easy to use tool to integrate validated interactions into transcriptional regulatory networks in plants and other species.

## Materials and Methods

### Validation of regulated TF targets in isolated plant cells

To identify the direct regulated targets of the 14 TFs in the ABA pathway that had both in planta ChIP (Song et al., 2016) and cis-binding motifs available (Weirauch et al., 2014), we expressed the TFs in isolated root cells using the TARGET system described in Brooks et al (Brooks et al., 2019) as follows. Arabidopsis Col-0 plants were grown in 1% w/v sucrose, 0.5 g per L MES, 0.5x MS basal salts (-CN), 2% agar, pH 5.7 for 10 days. Light conditions were 120 μmol m^-2^s^-1^ at constant temperature at 22°C, 16 h light, 8 h dark. Roots were harvested stirred with cellulase and macerozyme (Yakult, Japan) for 3 hours to remove the cell wall. Root protoplasts were filtered through 70 µm and then 40 µm cell strainers (BD Falcon, USA) and pelleted at 500 x g. Filtered root cells were washed with 15mL MMg buffer (400 mM mannitol, 10 mM MgCl_2_, 4mM MES pH 5.7) and resuspended to between 2-3 x 10^6^ cells per mL. Transfections of root cells were performed in a 50 mL conical tube by mixing 1 mL of root cell suspension with 120 μg of plasmid DNA, 1mL of PEG solution (40% polyethylene glycol 4000 (Millipore Sigma, USA), 400 mM mannitol, and 50 mM CaCl_2_) and vortexed gently for 5 seconds. After mixing, 50 mL of W5 buffer (154 mM NaCl, 125 mM CaCl2, 5 mM KCl, 5 mM MES, 5 mM glucose, pH 5.7) was added to the tube. Root cells were pelleted at 1,200 x g, and washed 3 times with W5 buffer. Cells transfected with a single TF in the RFP vector (pBOB11, available at https://gatewayvectors.vib.be/collection (Bargmann et al., 2013)) and another batch of cells transfected with a single TF in the GFP vector (pBOB11-GFP, available at https://gatewayvectors.vib.be/collection (Brooks et al., 2019)) were aliquoted into 3 replicate wells of a 24 well plate. The following day (18 hours) after TF expression and translation, transfected root protoplasts were treated with 35 µM CHX for 20 min before a 10 µM DEX treatment to induce TF nuclear import. Transfected root cells expressing the TF were sorted into GFP and RFP-expressing root cell populations by FACS 3 hours after DEX treatment.

To identify TF-regulated genes transcriptome analysis was performed. For this, cells expressing the candidate TF vs. EV were collected in triplicate and RNA-Seq libraries were prepared from their mRNA using the NEBNext® Ultra™ RNA Library Prep Kit for Illumina®. The RNA-Seq libraries were pooled and sequenced on the Illumina NextSeq 500 platform. The RNA-Seq reads were aligned to the TAIR10 genome assembly using HISAT2 (Kim et al., 2019) and gene expression estimated using the GenomicFeatures/GenomicAlignments packages (Lawrence et al., 2013). Gene counts were combined for each TF sample and the EV and differentially expressed genes in the TF transfected samples vs the EV samples were identified using the DESeq2 package (Love et al., 2014) with a TF+Batch model and an FDR adjusted p-value < 0.05. We filtered out genes that respond more than 5-fold to CHX treatment in transfected protoplasts (Brooks et al., 2019) from the lists of TF targets. Genes that are expressed in any of the protoplast experiments were used as the background for subsequent enrichment analyses in ConnecTF (Supplemental Table 9).

### TF-Target List Enrichment

Target list enrichment calculates the significance of the overlap between TF-targets in each queried TF analysis and each user-uploaded gene list. The p-values are calculated using the Fisher’s Exact Test adjusted with the Bonferroni correction. The background set of genes used for the calculation, which is by default all protein coding genes for both the Arabidopsis or maize instances of ConnecTF, can be manually set by the user by using the *Background Genes* option in the query page.

### Cis-motif Enrichment

Arabidopsis and maize cis-binding motif PWMs were collected from Cis-BP (Weirauch et al., 2014) (Build 2.0) and the 80 cis-motif clusters from Brooks et al. (Brooks et al., 2019) and converted to MEME format. The FIMO (Grant et al., 2011) tool within the MEME (Bailey et al., 2009) package was used to identify every occurrence of each cis-binding motif in the nuclear genome (i.e. excluding mitochondrial and chloroplast chromosomes) at a p-value < 0.0001 using the base frequency in the nuclear genome as the background model.

We chose to remove overlapping sites for the same cis-binding motifs, which are particularly common for repetitive motifs. For each cis-binding motif, when two sites overlap, the match with the lowest p-value is kept, and the other is removed until only non-overlapping matches remain. The number of matches for each cis-binding motif is tallied for each individual gene region, subdivided into 2000, 1000, and 500 bp upstream of transcription start site, the 5’ and 3’ untranslated regions (UTRs), coding sequence (CDS), intron, exon and the full region transcribed into mRNA (cDNA). If a match is found to be within a region shared by more than one gene, it is counted for all the genes that it is associated with.

To calculate enrichment of a cis-binding motif or cis-motif cluster for a particular individual TF within a given region in a target gene of a queried analysis, the Fisher’s Exact Test was used with a background of all individual cis-binding motifs or cis-motif clusters within that gene region, respectively. As in Target list enrichment, a user can upload a list of genes to use as the background, or use the default of all protein coding genes. The *P*-values are adjusted with the Bonferroni correction method.

If a Target Gene list (e.g. genes in a pathway of interest) is provided by the user, ConnecTF can also calculate the cis-binding motif enrichment for that gene list(s), separately. The p-values of motif enrichment on gene lists is adjusted with the Bonferroni correction as a group, independent of the correction performed on the queried analyses.

### Gene Set Enrichment

The Gene set enrichment tool calculates the significance of overlap between all possible pairwise combinations of target gene lists identified for any TF-targets queried. Significance of overlap is calculated using the Fisher’s exact test, using the default background of all protein coding genes, or the user uploaded background. Both the p-values for overlaps greater or equal to and lesser or equal to the one observed is calculated and displayed. All The p-values are then adjusted with the Bonferroni correction.

### Sungear

Sungear (Poultney et al., 2006) is a tool to display/visual overlaps between gene lists resulting from different queries, similar to a Venn diagram or UpSet plot (Lex et al., 2014). The vertices on the outer polygon are anchor points, vertices, containing gene lists for each TF-analysis queried. Circular nodes within the polygon represents gene sets that are in common between the indicated analyses. Each node has one or more arrows pointing to the vertices corresponding to the analyses which contains the genes. The gene sets exclusively found in that node represents the specific combination of analyses. The position of the node is approximately the midway point between the combination of analyses it represents.

In our implementation of Sungear, we enhanced the graph by calculating a p-value which indicates whether a node contains greater or fewer genes than expected given the total number of targets regulated by each of the queried analyses. Calculation was performed using the following method:

Let’s say there are n lists, each containing x_1_, x_2_ … x_n_ number of genes, with a total of x genes.

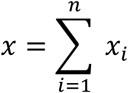

If a node *A*_1,2,4_ indicates genes that are exclusively in common with lists 1, 2, and 4. Then the expectation value, *e*, of a gene being in that node can be calculated from multiplying probability of being in the gene list and not being in the gene list respectively and *x*.

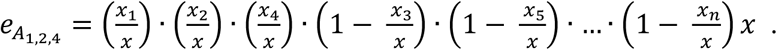

This will be a binomial distribution, where success is defined as the number of genes in the node A, and the failure is the number of genes not in node A (total genes - number of genes in node A). The p-value is calculated for each node by comparing the observed value to the expected value using the binomial test and adjusted using the Bonferroni correction.

### Code Availability

The source code including instructions for setting up a public or private instance of ConnecTF is available at https://github.com/coruzzilab/connectf_server.

### Data Availability

All raw sequencing data from this project have been deposited in the Gene Expression Omnibus (GEO) database, https://www.ncbi.nlm.nih.gov/geo (accession no. GSE152405).

## Acknowledgements

We would like to acknowledge Reetu Tuteja for the contributions she made in the early stages of this project. We would like to thank Lauriebeth Leonelli for vital advice on the design of the site and for suggesting the name ConnecTF. Finally, we would like to thank Dennis Shasha, Carol Huang and members of the Coruzzi Lab for feedback throughout the development of ConnecTF, in particular Chia-Yi Cheng, Gil Eshel, and Joseph Swift. This work was supported by NIH Grant GM032877 and NSF-PGRP: IOS-1339362 to G.C., NIH NIGMS Fellowship F32GM116347 to M.D.B., and a Plant Genomics Grant from the Zegar Family Foundation (A160051).

