## Supplemental Figures and Data File for "ConnecTF: A platform to build gene networks by integrating transcription factor-target gene interactions"

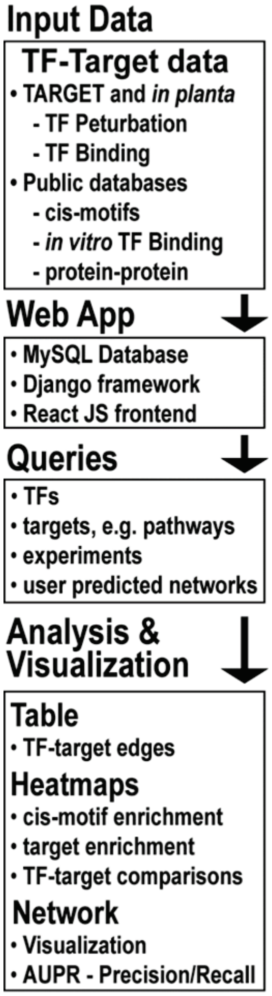


**Supplemental Figure 1 - Diagram of ConnecTF from data to analysis tools.** ConnecTF can store multiple types of TF-target interaction data including but not limited to TF-perturbation, TF-binding, in vitro binding (e.g. DAP-seq (O'Malley et al., 2016)), cis-binding motifs, protein-protein interaction and open chromatin (Input Data). The backend and frontend of the ConnecTF platform is species-independent and built using common software such as MySQL, Django and React JS (Web App). To make the tool accessible to both researchers new to the ConnecTF site and for those who wish to build complex queries for parsing multiple datasets for the many available TFs, queries can be built using the graphical interface or by typing into the search text box (Queries). Many tools are built into the site to enable further analysis of the results returned by the query, such as network visualization, predicted network refinement/pruning, cis-motif enrichment, target list enrichment, pairwise TF-target list comparisons and common/shared TF-target list comparisons (see Figure 1).

| | |
|---|---|
| AUPR | 0.2138 |
| AUPR Random | 0.1892 |
| p-value | <0.001 |
| Precision Threshold | 0.25 |
| Edge Score Threshold | |
| Number of edges | 24,405/1,000,000 |
| Number of TFs | 1,623/1,742 |
| Number of Targets | 9,831/22,672 |

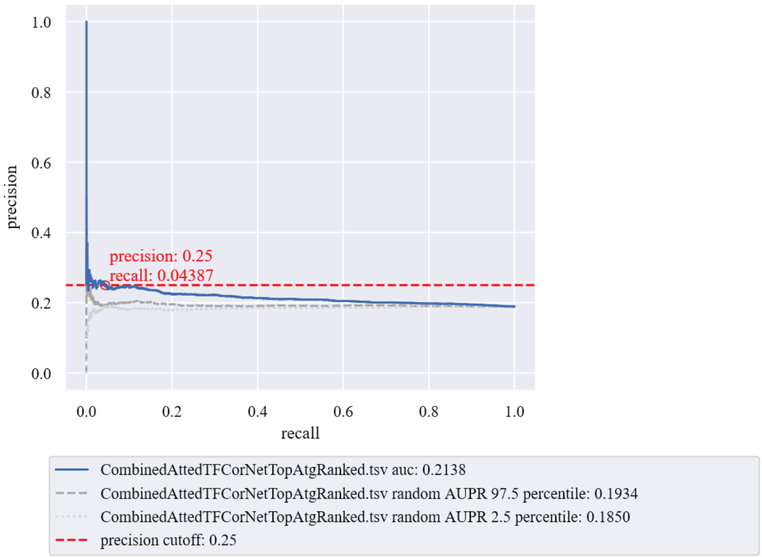


**Supplemental Figure 2 – Case Study 2: Precision/recall analysis on the Atted-II co-expression network.** This screenshot from the *Network* tab shows how we used ConnecTF to perform precision/recall analysis on the top scoring 1 million predicted interactions between 1,742 Arabidopsis TFs and 22,672 target genes in the co-expression GRN downloaded from the Atted-II database (v9.2) (Obayashi et al., 2018). Validated TF-regulation datasets housed in ConnecTF were used as “gold standard” edges for the network pruning/refinement.

| | NLP7 regulated/not bound (505) |
|---|---|
| AT1G72830 (NF-YA3) (1889) | 7.10e-65 (150) |
| AT3G49940 (LBD38) (1387) | 4.75e-61 (127) |
| AT5G67420 (LBD37) (2670) | 1.41e-59 (169) |
| AT5G67420 (LBD37) (1524) | 2.60e-50 (120) |
| AT2G22430 (HB6) (2566) | 1.25e-42 (143) |
| AT2G39250 (SNZ) (1070) | 5.06e-28 (76) |
| AT5G62430 (CDF1) (2656) | 1.13e-22 (113) |
| AT5G62430 (CDF1) (3568) | 2.63e-22 (134) |
| AT5G04340 (ZAT6) (3128) | 1.96e-20 (120) |
| AT2G33550 (ASR3) (2003) | 8.79e-20 (91) |
| AT3G49940 (LBD38) (2301) | 2.15e-18 (96) |
| AT4G39780 (ERF060) (2219) | 2.07e-15 (88) |
| AT5G05410 (DREB2A) (957) | 1.36e-10 (46) |
| AT4G39780 (ERF060) (523) | 1.11e-8 (30) |

**Supplemental Figure 3 – Case Study 3: Enrichment of TF2 targets with NLP7 indirect targets reveals influential downstream TFs.** A screenshot from ConnecTF showing the results of using the *Target Enrichment* tool. We found that the direct regulated targets identified in root cells for all 8 $TF_2$s are enriched for NLP7 indirect targets. This analysis reveals that NF-YA3, LBD37 and LBD38 are particularly enriched in NLP7 indirect targets (Fisher's exact test).

**Supplemental Data File- ConnecTF queries used to generate figures and tables**

**Figure 2**

| | |
|---|---|
| *Query*: | all_expression |
| *Target Genes*: | Abscisic_Acid_Responsive |
| *Filter TFs*: | ABA TFs |
| *Background*: | TARGET_Expressed |
| Notes: | all_Expression is shorthand for all_tfs[Experiment_Type=Expression] |

Explanation: This query returns any analysis that has expression as the "Experiment_Type" in the metadata (e.g. RNA-seq or microarray experiments). The targets are limited to the ABA-responsive genes from Song et al. (Song et al., 2016) and the TFs are filtered for the 14 TFs in ABA signaling. The background used for enrichment analyses are only genes expressed in the TARGET assays from this study and Brooks et al. (Brooks et al., 2019).

**Figure 3**

| | |
|---|---|
| *Query*: | AT2G46680[log2fc<0] or AT2G46680[log2fc>0] or AT1G22640[log2fc<0] or AT1G22640[log2fc>0] or AT5G04340[log2fc<0] or AT5G04340[log2fc>0] |
| *Background*: | TARGET_Expressed |

Explanation: This query demonstrates how ConnecTF can be used to parse the TF-targets of one or more experiments. In this example, the targets of three TFs (HB7, MYB3 and ZAT6) into the induced and repressed targets of each TF. As log2 fold-change values are only associated with TF-regulation experiments in ConnecTF, it also acts as a filter to exclude TF-binding experiments. The background used for enrichment analyses are only genes expressed in the TARGET assays from this study and Brooks et al. (Brooks et al., 2019).

**Figure 4**

*Query*:
1. AT2G46680[log2fc<0] or
AT2G46680[log2fc>0] or
AT2G46680[EDGE_TYPE='in planta:Bound']
2. AT1G22640[log2fc<0] or
AT1G22640[log2fc>0] or
AT1G22640[EDGE_TYPE='in planta:Bound']
3. AT5G04340[log2fc<0] or
AT5G04340[log2fc>0] or
AT5G04340[EDGE_TYPE='in planta:Bound']

*Background*: 1-3. TARGET_Expressed

Explanation: This query again parses the TF-targets of the three TFs, HB7, MYB3 and ZAT6 into induced vs. repressed TF-regulated and TF-bound targets. The background used for enrichment analyses are only genes expressed in the TARGET assays from this study and Brooks et al. (Brooks et al., 2019).

**Figure 5**

*Query*: A:(AT5G04340[log2fc>0] and
AT5G04340[EDGE_TYPE='in planta:Bound']) or
(AT5G04340[log2fc<0] and
AT5G04340[EDGE_TYPE='in planta:Bound'])

*Background*: TARGET_Expressed

Explanation: In this query, grouping of filters enables the user to find the overlap between targets for a single TF from multiple experiments, in this example targets that are induced *and* TF-bound and targets that are repressed *and* TF-bound. The targets are limited to the ABA-responsive genes from Song et al. (Song et al., 2016). The background used for enrichment analyses are only genes expressed in the TARGET assays from this study and Brooks et al. (Brooks et al., 2019).

**Figure 6**

*Query*: all_expression[TISSUE/SAMPLE='Root Protoplasts']
*Target Network*: Root_Nitrogen_Predicted_Network
*Background*: TARGET_Expressed
Notes: all_Expression is shorthand for all_tfs[Experiment_Type=Expression]

Explanation: This query was used to demonstrate the precision/recall functionality of ConnecTF. A user can upload an inferred network with TF-target gene interactions in ranked order as a *Target Network* on the query page. In this example, the inferred network is from the time-series nitrogen response in Arabidopsis roots (Brooks et al., 2019). A precision/recall analysis will be performed automatically in the *Network* tab (Note: if an unranked network is uploaded, the precision/recall analysis will be meaningless). The background used for enrichment analyses are only genes expressed in the TARGET assays from this study and Brooks et al. (Brooks et al., 2019).

**Figure 7**

*Query*:
1. AT4G24020[EXPERIMENT_TYPE=Expression] and AT4G24020[TECHNOLOGY/METHOD=ChIPseq]
2. AT4G24020[EXPERIMENT_TYPE=Expression] and not AT4G24020[TECHNOLOGY/METHOD=ChIPseq]
3. all_expression[TISSUE/SAMPLE='Root Protoplasts']

*Filter TFs*:
1. None
2. None
3. Targets from query 1 e.g. Bound and regulated (direct) by NLP7

*Target Genes*:
1. None
2. None
3. Targets from query 2 e.g. Regulated but unbound (indirect) by NLP7

Notes: all_Expression is shorthand for all_tfs[Experiment_Type=Expression]

Explanation: This set of three queries demonstrates how ConnecTF can be used in an iterative fashion, wherein the results from the first two queries are saved and as inputs for the *Filter TFs* and *Target Genes* fields and act as filters in the third query. We demonstrate this functionality by using ConnecTF to perform Network Walking (Brooks et al., 2019), and chart a network path from NLP7 (the $TF_1$), via the intermediate $TF_2$s which it directly regulates (query 1 results), to its indirect targets (query 2 results).

**Supplemental Table 2**

| | |
|---|---|
| *Query*: | 1. AT4G24020 |
| | 2. AT4G24020 |
| *Target Genes*: | 1. None |
| | 2. Nitrogen_by_Time |

Explanation: These examples demonstrate the most basic functionality of ConnecTF. That is 1) a query to return all the experiments available for a TF, and 2) to interrogate how a TF regulates a target gene list(s) of interest. In this example, the first query returns the seven datasets in ConnecTF for the nitrogen master regulator NLP7. The second query, we limit the targets to those responding to nitrogen by time in either roots or shoots by selecting the Target Gene List.

**Supplemental Table 4**

| | |
|---|---|
| *Query*: | 1. all_tfs[log2fc>0] |
| | 2. all_tfs[log2fc<0] |
| *Target Genes*: | 1. Abscisic_Acid_Responsive |
| | 2. Abscisic_Acid_Responsive |
| *Filter TFs*: | 1. ABA TFs |
| | 2. ABA TFs |
| *Background*: | 1. TARGET_Expressed |
| | 2. TARGET_Expressed |

Explanation: This pair of queries returns expression experiments (RNA-seq or microarray) split into induced and repressed targets. The targets are limited to the ABA-responsive genes from Song et al. (Song et al., 2016) and the TFs are filtered for the 14 TFs in ABA signaling. This was used to compare the TF-induced/repressed targets of the 14 TFs to genes that were induced/repressed in response to the ABA signal. The background used for enrichment analyses are only genes expressed in the TARGET assays from this study and Brooks et al. (Brooks et al., 2019).

**Supplemental Table 5**

*Query*: expand("$filter_tf[log2fc>0] or $filter_tf[log2fc<0]",or)
*Filter TFs*: ABA TFs
*Background*: TARGET_Expressed

Explanation: This query returns the induced and repressed targets separately for each of the 14 ABA responsive TFs. It utilizes a special “expand” function that has been added to the query system of ConnecTF and enables users to write a query that will repeat a more complex query for each TF in the Filter TFs list, replacing “$filter_tf” with each of the TFs in the list and connecting them with “and” or “or” depending on the last argument in the function. For example, this query could also be written as:
(AT5G67300 [log2fc<0] or AT5G67300 [log2fc>0]) or
(AT1G22640 [log2fc<0] or AT1G22640 [log2fc>0]) or
(AT1G49720 [log2fc<0] or AT1G49720 [log2fc>0]) or
(AT1G51140 [log2fc<0] or AT1G51140 [log2fc>0]) or
(AT2G22430 [log2fc<0] or AT2G22430 [log2fc>0]) or
(AT2G46270 [log2fc<0] or AT2G46270 [log2fc>0]) or
(AT2G46680 [log2fc<0] or AT2G46680 [log2fc>0]) or
(AT4G01120 [log2fc<0] or AT4G01120 [log2fc>0]) or
(AT4G34000 [log2fc<0] or AT4G34000 [log2fc>0]) or
(AT4G37790 [log2fc<0] or AT4G37790 [log2fc>0]) or
(AT5G04340 [log2fc<0] or AT5G04340 [log2fc>0]) or
(AT5G04760 [log2fc<0] or AT5G04760 [log2fc>0]) or
(AT5G05410 [log2fc<0] or AT5G05410 [log2fc>0]) or
(AT5G43840 [log2fc<0] or AT5G43840 [log2fc>0])

The background used for enrichment analyses are only genes expressed in the TARGET assays from this study and Brooks et al. (Brooks et al., 2019).

**Supplemental Table 6**

*Query*: all_tfs[log2fc<0] or all_tfs[log2fc>0] or all_dap
*Filter TFs*: ABA TFs
*Background*: TARGET_Expressed
Notes: all_dap is shorthand for all_tfs[Edge_type=’in vitro:Bound:DAP’ or Edge_type=’in vitro:Bound:ampDAP’]

Explanation: In order to compare the overlap between induced (or repressed) and in vitro TF-bound targets this query and the *Gene Set Enrichment* tool in ConnecTF were used. The TFs are filtered for the 14 TFs in ABA signaling. The background used for enrichment analyses are only genes expressed in the TARGET assays from this study and Brooks et al. (Brooks et al., 2019). The results from *Gene Set Enrichment* were exported to a csv file. This file contains every pairwise comparison between the 14 TFs, and therefore only the rows corresponding to comparisons between bound and regulated targets for a single TF were retained while the remaining rows were discarded.

**Supplemental Table 6**

| | |
|---|---|
| *Query*: | all_tfs[log2fc<0] or all_tfs[log2fc>0] or in_planta_bound |
| *Filter TFs*: | ABA TFs |
| *Background*: | TARGET_Expressed |
| Notes: | in_planta_bound is shorthand for all_tfs[Edge_type='in planta:Bound'] |

Explanation: In order to compare the overlap between induced (or repressed) and TF-bound targets this query and the *Gene Set Enrichment* tool in ConnecTF were used. The TFs are filtered for the 14 TFs in ABA signaling. The background used for enrichment analyses are only genes expressed in the TARGET assays from this study and Brooks et al. (Brooks et al., 2019). The results from *Gene Set Enrichment* were exported to a csv file. This file contains every pairwise comparison between the 14 TFs, and therefore only the rows corresponding to comparisons between bound and regulated targets for a single TF were retained while the remaining rows were discarded.

**Supplemental Table 8**

*Query*: expand("($filter_tf[Edge_type='in planta:Bound'] and $filter_tf[log2fc>0]) or ($filter_tf[Edge_type='in planta:Bound'] and $filter_tf[log2fc<0])",or)

*Filter TFs*: ABA TFs

*Background*: TARGET_Expressed

Explanation: This query was used to identify enriched cis-motif clusters in the TF-target genes that are direct regulated (induced) *and* TF-bound in vivo (ChIP-seq), or direct regulated (repressed) and TF-bound in vivo (ChIP-seq) . The query utilizes a special "expand" function that has been added to the query system of ConnecTF and enables users to write a query that will repeat a more complex query for each TF in the Filter TFs list, replacing "$filter_tf" with each of the TFs in the list and connecting them with "and" or "or" depending on the last argument in the function. For example this query could also be written as:

((AT5G67300 [Edge_type='in planta:Bound'] and AT5G67300 [log2fc>0]) or
(AT5G67300 [Edge_type='in planta:Bound'] and AT5G67300 [log2fc<0])) or
((AT1G22640 [Edge_type='in planta:Bound'] and AT1G22640 [log2fc>0]) or
(AT1G22640 [Edge_type='in planta:Bound'] and AT1G22640 [log2fc<0])) or
((AT1G49720 [Edge_type='in planta:Bound'] and AT1G49720 [log2fc>0]) or
(AT1G49720 [Edge_type='in planta:Bound'] and AT1G49720 [log2fc<0])) or
((AT1G51140 [Edge_type='in planta:Bound'] and AT1G51140 [log2fc>0]) or
(AT1G51140 [Edge_type='in planta:Bound'] and AT1G51140 [log2fc<0])) or
((AT2G22430 [Edge_type='in planta:Bound'] and AT2G22430 [log2fc>0]) or
(AT2G22430 [Edge_type='in planta:Bound'] and AT2G22430 [log2fc<0])) or
((AT2G46270 [Edge_type='in planta:Bound'] and AT2G46270 [log2fc>0]) or
(AT2G46270 [Edge_type='in planta:Bound'] and AT2G46270 [log2fc<0])) or
((AT2G46680 [Edge_type='in planta:Bound'] and AT2G46680 [log2fc>0]) or
(AT2G46680 [Edge_type='in planta:Bound'] and AT2G46680 [log2fc<0])) or
((AT4G01120 [Edge_type='in planta:Bound'] and AT4G01120 [log2fc>0]) or
(AT4G01120 [Edge_type='in planta:Bound'] and AT4G01120 [log2fc<0])) or
((AT4G34000 [Edge_type='in planta:Bound'] and AT4G34000 [log2fc>0]) or
(AT4G34000 [Edge_type='in planta:Bound'] and AT4G34000 [log2fc<0])) or
((AT4G37790 [Edge_type='in planta:Bound'] and AT4G37790 [log2fc>0]) or
(AT4G37790 [Edge_type='in planta:Bound'] and AT4G37790 [log2fc<0])) or
((AT5G04340 [Edge_type='in planta:Bound'] and AT5G04340 [log2fc>0]) or
(AT5G04340 [Edge_type='in planta:Bound'] and AT5G04340 [log2fc<0])) or
((AT5G04760 [Edge_type='in planta:Bound'] and AT5G04760 [log2fc>0]) or
(AT5G04760 [Edge_type='in planta:Bound'] and AT5G04760 [log2fc<0])) or
((AT5G05410 [Edge_type='in planta:Bound'] and AT5G05410 [log2fc>0]) or
(AT5G05410 [Edge_type='in planta:Bound'] and AT5G05410 [log2fc<0])) or
((AT5G43840 [Edge_type='in planta:Bound'] and AT5G43840 [log2fc>0]) or
(AT5G43840 [Edge_type='in planta:Bound'] and AT5G43840 [log2fc<0]))

The TFs are filtered for the 14 TFs in ABA signaling. The background used for enrichment analyses are only genes expressed in the TARGET assays from this study and Brooks et al. (Brooks et al., 2019).

**Supplemental Figure 2**

*Query*: all_expression
*Target Network*: Atted-II Co-expression Network
Notes: all_Expression is shorthand for all_tfs[Experiment_Type=Expression]

Explanation: This query was used to demonstrate the precision/recall functionality of ConnecTF. A user can upload an inferred network with TF-target gene interactions in ranked order as a *Target Network* on the query page. In this example, the inferred network is the top scoring 1 million predicted interactions between 1,742 Arabidopsis TFs and 22,672 target genes in the co-expression GRN downloaded from the Atted-II database (v9.2) (Obayashi et al., 2018). A precision/recall analysis will be performed automatically in the *Network* tab (Note: if an unranked network is uploaded, the precision/recall analysis will be meaningless).

**Supplemental Figure 3**

Query: all_expression[TISSUE/SAMPLE='Root Protoplasts']
Filter TFs: Targets bound and regulated (direct targets) by NLP7 (Query 1 for Figure 7)
Target List: Genes regulated but unbound (indirect targets) by NLP7 (Query 2 for Figure 7)

Explanation: This is the same query as that #3 from Figure 7. The results are viewed in the *Target List Enrichment* tab to identify the $TF_2$s that are direct targets of NLP7 that are most influential, most specific and most enriched for regulation of NLP7 indirect targets.
